# NK cells undergo transcriptional and functional reprogramming following *Streptococcus pneumoniae* infection

**DOI:** 10.1101/2025.09.19.677332

**Authors:** Júlia Torné, Claudia Chica, Tiphaine Camarasa, Bernd Jagla, Matilde Diaz Enes, Aymeric Zellner, Sébastien Mella, Valentina Libri, Mélanie Anne Hamon

**Affiliations:** Institut Pasteur, Université Paris Cité, Chromatin and Infection Laboratory, F-75015, Paris, France; Institut Pasteur, Université Paris Cité, Bioinformatics and Biostatistics Hub, F-75015 Paris, France; Institut Pasteur, Université Paris Cité, Single Cell Biomarkers UTechS, F-75015 Paris, France; Current address: Infection Immunology laboratory, Department of Biomedicine, University of Basel, Basel, Switzerland; Current address : Institut Pasteur, ILC development and inflammation Laboratory, Paris, France

## Abstract

Natural Killer (NK) cells are cytotoxic lymphocytes and key mediators of innate immunity, essential for combating viral infections and cancer. Notably, they exhibit immunological memory, generating a stronger response upon re-exposure to the same stimulus. While NK cell memory holds promise for infection control, its role in bacterial infections remains poorly understood. Previously, we demonstrated that *Streptococcus pneumoniae* induces long-term, specific, and protective NK cell memory. In this study, we performed single-cell RNA-seq to uncover how NK cells respond to *S. pneumoniae* infection. Our findings reveal that challenged Memory (cMemory) NK cells undergo transcriptional reprogramming following *S. pneumoniae* infection and have a differential transcriptional response upon reinfection. In addition, we identified distinct cMemory NK cell subpopulations, with responding cMemory NK cells displaying a general enhanced activation, proliferation, and cytotoxic activity. These findings support a novel role for NK cells in the context of bacterial infections, thereby opening avenues for harnessing the potential of innate immune memory for therapeutic applications.

## INTRODUCTION

The identification of memory responses driven by innate immune cells has challenged the long-standing belief that only adaptive immune cells can recall prior exposures. The ability of innate immune cells, such as macrophages and monocytes, to display memory properties is the subject of intense research and is referred to as trained immunity. This form of innate immune memory provides enhanced resistance to reinfection in a non-specific manner, and is accompanied by the acquisition of permanent epigenetic and metabolic changes (Netea et al., 2011; Quintin et al., 2014; Saeed et al., 2014; Divangahi et al., 2021).

Natural Killer (NK) cells are cytotoxic lymphocytes, important mediators of innate immunity and well characterised for their role in fighting viral infections and cancer. Traditionally, NK cells have been described as short-lived first responders of the innate immune system. However, recent studies are revealing new adaptive NK cell properties including cell residency and the acquisition of memory in an antigen-specific manner, providing an enhanced immune response upon re-exposure to the same original stimulus, in contrast to trained immunity which is unspecific (Cerwenka & Lanier, 2016; Geary & Sun, 2017; O’Sullivan et al., 2015). NK cell memory has been extensively studied in the context of viral infections, where specific subpopulations of NK cells are responsible for recognising viral components, which trigger their clonal expansion and long-term persistence (Lopez-Vergès et al., 2011; Sun et al., 2009). This results in the development of a virus-specific subpopulation of NK cells with unique and lasting chromatin accessibility states, promoting the expression of key genes to enable Memory NK cells to mount a more robust immune response when re-exposed to the same virus (Lau et al., 2018).

In contrast to responses to viruses, the role of NK cell memory in bacterial infections has been poorly studied and remains very descriptive. Studies on *Ehrlichia muris* showed that NK cells from infected mice could protect against a subsequent lethal infection by the same bacterium (Habib et al., 2016; Thirumalapura et al., 2008). Another study on BCG vaccination pointed to their ability to show memory responses, since transfer of memory-like NK cells primed by BCG inoculation conferred protection to recipient mice against *M. tuberculosis* infection by reducing bacterial counts in the lungs at 30 days post-infection (Venkatasubramanian et al., 2017). In a recent study, we showed that NK cells developed specific and long-term memory properties to *S. pneumoniae,* characterised by a protection against lethal infection upon transfer into naïve mice, and in a manner that is distinct from viral-induced NK cell memory (Camarasa et al., 2023). Memory NK cells conferred protection and significantly reduced the bacterial burden for at least 12 weeks, in a pathogen specific manner. Interestingly, although *S. pneumoniae* is mainly extracellular, when placed *in vitro*, NK cells directly sense and respond to this bacterium. The observed specificity of NK cell memory to *S. pneumoniae* supports the antigen-specific memory characteristics of NK cells during bacterial infection, distinguishing this response from trained immunity. Although memory features of NK cells were described phenotypically, the identity and the transcriptional specificities for acquisition and maintenance of memory to a bacterial infection are unknown.

Importantly, growing evidence indicates that NK cells are not an homogeneous population, they exist in subpopulations defined by unique transcriptional landscapes that are thought to reveal not only their origin and localisation, but also their specific effector roles (Crinier et al., 2018; Schuster et al., 2023a). Within this landscape, Memory NK cells have been defined as a subpopulation of NK cells that, when activated, give rise to an expanded subpopulation, characterised by an enhanced response (Panjwani et al., 2024; Rückert et al., 2022). The study of NK cell subpopulations is now essential for defining memory NK cells to bacterial infections and for a better understanding of NK cell memory origin and responses.

In this study we perform a comprehensive single cell transcriptional analysis of challenged Naïve and Memory (cNaïve and cMemory) murine NK cells in response to an intranasal infection with *S. pneumoniae*. We reveal that NK cells are transcriptionally reprogrammed following a first infection and new Memory populations arise. We demonstrate that response to a secondary infection is different from the primary infection, with enhanced expression of genes related to NK cell activation, residency, survival, and cytotoxic activity. We further validate our findings at the protein level following *S. pneumoniae* challenge and show that enhanced proliferation and cytotoxic response are intrinsic properties of Memory NK cells. These findings therefore establish that bacterial infection drives the establishment of new NK cell populations and shed light into the features that characterise the enhanced functionality of Memory NK cells.

## RESULTS

### 1. *S. pneumoniae* infection drives significant changes in multiple NK cell clusters

Our recent discovery that bacterial infection can induce a protective NK cell memory phenotype led us to investigate the underlying NK cell memory responses responsible for this protection. We hypothesised that NK cells undergo transcriptional reprogramming after a first infection, enabling them to mount a different transcriptional program during a subsequent infection. To address this question, we used a model of adoptive transfer of NK cells (**Figure 1A**) (Camarasa et al., 2023). CD45.2 mice were infected intranasally with a sub-lethal dose of *S. pneumoniae* (SPN) (5 x 10^5^ CFU/ml) or PBS in the control group as previously described. 21 days later, when the infection was completely cleared and NK cells were back to their basal state (Camarasa et al., 2023), D21PBS NK cells (Naïve, from the control mice) and D21SPN NK cells (Memory, from the infected mice) were highly purified by negative selection from spleens (**Supplementary** Figure 1A) and transferred intravenously by retro-orbital injection into CD45.1 recipient mice. Recipient mice that had received either Naïve or Memory NK cells were infected with a challenge of *S. pneumoniae* (SPN; 1 × 10⁷ CFU/mL) 24 hours post-transfer. 40 hours post-infection, lungs were harvested, and CD45.2+ transferred NK cells were sorted (gating strategy shown in **Supplementary** Figure 1B). As expected, the transfer of Memory NK cells conferred a protective effect to the recipient mice, evidenced by reduced bacterial counts in their lungs (**Supplementary** Figure 1C). Importantly, we have previously shown that protection by memory NK cells does not imply an increased recruitment of other immune cells nor an increased inflammatory response in the lung (Camarasa et al., 2023). To investigate the intrinsic transcriptional responses of challenged transferred cells, we performed single cell RNA sequencing (scRNA-seq) on transferred and challenged NK cells (**Figure 1A**). The challenged Naïve (cNaïve) cells were encountering the infection for the first time, while the challenged Memory (cMemory) NK cells were responding to a re-infection. To assess the heterogeneity of NK cell responses to infection in the lung, we performed an unsupervised droplet-based single-cell mRNA sequencing analysis using the 10X Genomics Chromium technology. We analysed transferred (CD45.2) cMemory and cNaïve NK cells. After quality control, 4918 (2453 Naïve and 2465 Memory) mouse NK cells were normalised, integrated, and clustered. We explored a range of clustering resolutions that, as expected, affected the cluster sizes but had no effect on the distribution of cNaïve and cMemory cells among clusters (Supplementary Figure 1D). We opted for the lowest resolution that would lead to the identification of clusters with interpretable transcriptional profiles. We thus defined 6 clusters that describe distinct cellular states of the NK cell population under study and include both cNaïve and cMemory cells: Cluster 0 contained 1886 cells, Cluster 1 contained 1339 cells, Cluster 2 contained 729 cells, Cluster 3 contained 381, Cluster 4 contained 258 cells and Custer 5 contained 225 cells (**Figure 1B** and **Figure 1D**).

**Figure 1.**
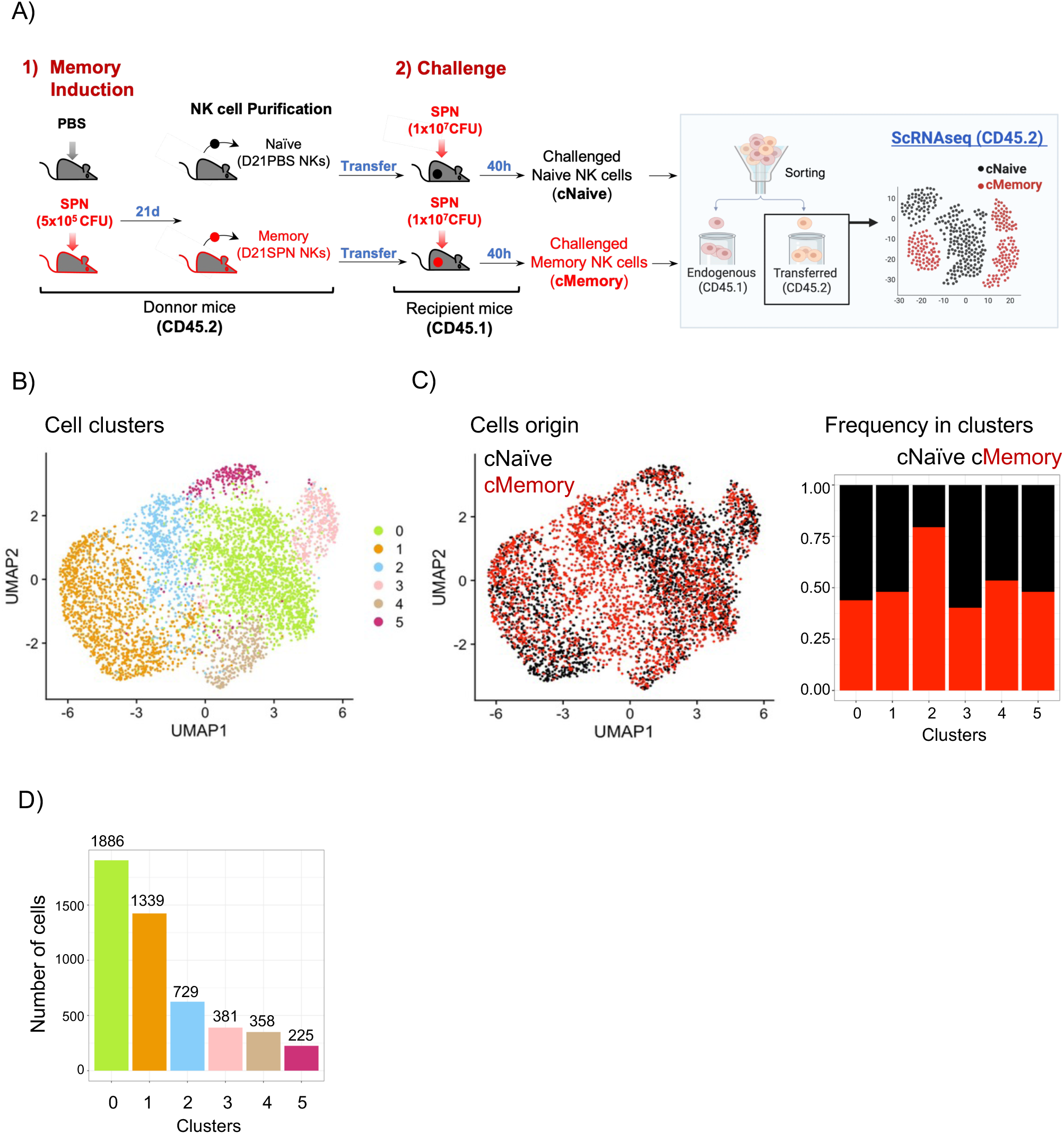
scRNA-seq reveals different NK cell clusters in response to *S. pneumoniae.* **A)** Experimental scheme. C57BL/6 mice (donor mice, CD45.2+) were intranasally infected with either PBS or a sub-lethal dose of *S. pneumoniae* (SPN, 5x10^5^ CFU) for two consecutive days. After 21 days, NK cells from spleens were highly purified (D21PBS or D21SPN NKs) and 5x10^5^ cells were transferred intravenously into Ly5.1 Naïve mice (recipient mice, CD45.1+). One day after, all recipient mice were intranasally infected with a lethal dose of *S. pneumoniae* (1x10^7^ CFU). Lungs were collected 40h post-infection for the sorting of the transferred cells (CD45.2+). **B)** Two-dimensional UMAP representation of 2453 cNaïve and 2465 cMemory NK cells colored by clusters calculated according to their transcriptional resemblance. **C)** UMAP representation of cells colored by origin, cNaïve (black) or cMemory (red) (**left panel).** Proportion of cNaïve (black) and cMemory (red) cells per cluster (**right panel**).

The existence of different subpopulations indicates that transferred NK cells are generally a heterogeneous population of cells. A different representation of the clusters by their origin, shows that most of the clusters are composed of a mix of cMemory and cNaïve NK cells (**Figure 1C**). However, one cluster is mainly composed of cMemory NK cells. Indeed, cluster 2 is dominated by cMemory NK cells, representing 80% of the total NK cells comprised in this cluster. These results indicate that NK cells exhibit a significantly different transcriptional response to infection depending on whether they had previously been exposed to bacteria or not.

### 2. Mixed NK cell clusters display functional diversity

To understand the identity and function of each cluster — most of which contain a mix of cMemory and cNaïve NK cells — we focused on the features shared by cells within each individual cluster. First, we report the top marker genes of the three biggest Clusters: 0, 1 and 2 (**Figure 2A**). Additionally, we highlight significant marker genes (**Figure 2B**), and classify top and selected marker genes by genes encoding for secreted proteins, cell surface markers, and transcription factors, which emphasised the individual characteristics of every cluster (**Figure 2C**). In parallel to the marker genes detection, to distinguish the most important biological processes ongoing in every cluster, we performed competitive gene set tests, providing the normalised enrichment score (NES) as the primary statistic for examining gene sets enriched in each cluster (**Figure 2D and Supplementary Table 5**).

**Figure 2.**
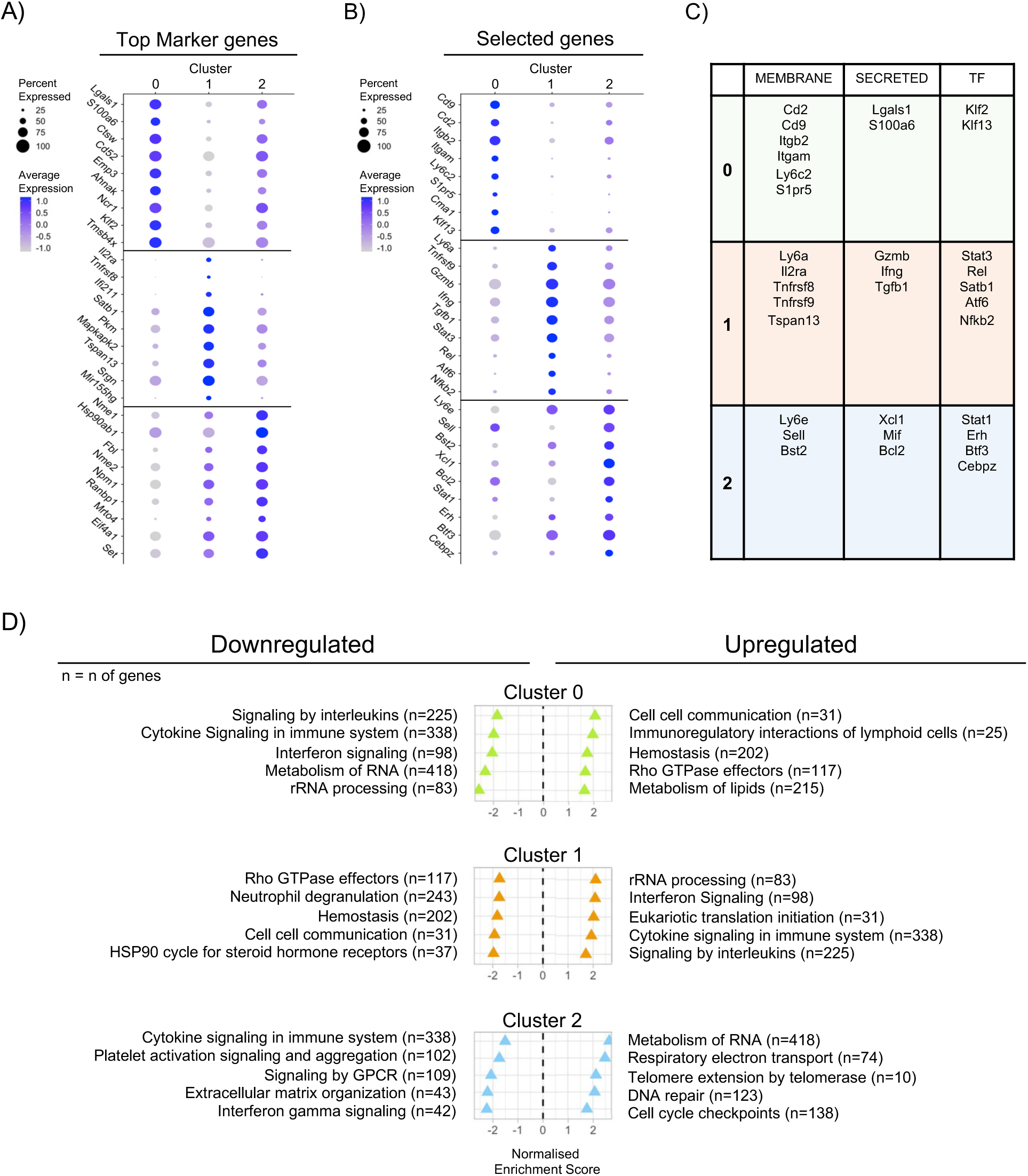
Characterizing the functional diversity of NK cell clusters. **A)** Dot plot representing the expression of top 9 marker genes, conserved among cNaïve and cMemory cells and ranked by their average log fold change, for clusters 0, 1 and 2. The size of the dots represents the percentage of cells expressing the corresponding gene and the colour represents its average expression. **B)** Dot plot representing the expression of selected relevant marker genes for clusters 0, 1 and 2. The size of the dots represents the percentage of cells expressing the corresponding gene and the colour represents its average expression. **C)** Classification of marker genes per cluster (1^st^ column) into membrane markers, secreted proteins, or transcription factors (TF). **D)** Gene Set Enrichment Analysis (GSEA) for each set of markers conserved among cNaïve and cMemory in every cluster. Selected REACTOME gene sets significantly enriched (False Discovery rate < 0.05) among upregulated or downregulated genes in clusters 0, 1 and 2 are compared using the Normalized Enrichment Score (NES). n = number of genes in every gene set.

Cluster 0 is defined by genes that are classically expressed in steady-state NK cells, suggesting that these cells are not actively responding to infection (Crinier et al., 2018). These include *Cd2* and *Cd9*, coding for surface proteins involved in immune cell interactions, *Ncr1*, *NKp46*, *Ly6c2*, *Itgam* (Cd11b), coding for cell surface receptors, *Lgals1*, coding for a galectin, and *Klf2* and *Klf13*, two transcription factors important for NK cell proliferation, survival, and migration and for cytotoxicity and cytokine production, respectively. Functional analysis displayed an enrichment of gene sets related to cell communication, immunoregulatory interactions, or hemostasis along with downregulation of gene sets linked to inflammation and immune response, supporting the notion that these cells are not responding to infection.

In contrast, cells in Cluster 1 show upregulation of *Il2ra, Ly6a*, and *Tnfrsf8* encoding surface markers linked to NK cell activation, as well as *Ifng* and *Gzmb*, coding for secreted proteins that perform NK cell effector functions. This cluster also exhibits a specific set of transcription factors (TF) suggesting intense chromatin remodelling, including *Stat3*, involved in various NK cell response functions, and *Nffib2* and *Rel* encoding for proteins involved in the NF-κB signalling pathway. The expressed marker genes of this cluster strongly suggest that these cells are actively responding to infection. Indeed, functional analysis reveals enrichment in interferon and cytokine signalling, interleukin signalling and active transcription and translation, in line with an active immune response.

In Cluster 2, in which 80% of cells are cMemory NK cells, we observe expression of specific surface markers such as *Ly6e, Sell* or *Bts2*. Additionally, a different set of secreted molecules like *Xcl1 or Mif,* both important for modulating NK cell recruitment and immune response, are specifically expressed in this cluster. Furthermore, expression of *Bcl2* is upregulated, encoding for an anti-apoptotic protein. Finally, cells in Cluster 2 express a different set of transcription factors, suggesting distinct chromatin dynamics. These include *Stat1, Erh, Btf3*, or *Cebpz*, all involved in regulating various aspects of NK cell biology, including activation, cytokine production, cytotoxicity. Functional analysis points to increased transcription and translation as well as telomere extension, DNA repair, and cell cycle checkpoints. However, pathways related to active immune responses and genes related to NK cell response (GzmB, Ifng,,..) are downregulated, suggesting that cMemory NK cells in this cluster are not actively responding to the infection, similarly to Cluster 0, but with a different transcriptional program.

Clusters 3, 4 and 5 are smaller clusters, representing only 19% of analysed cells. However, they are also characterised by specific top marker genes that diverge from the ones in clusters 0, 1 or 2 (**Supplementary** Figure 2A and 2B). Cluster 3 is primarily defined by its top two marker genes, *Hspa1a* and *Hspa1b* encoding for heat shock response genes. Cluster 4 is characterised by high expression of *Ccl3* and *Ccl4*, encoding for chemokines involved in NK cell migration and response to infection. Other marker genes of Cluster 4 include TF such as *Nffibid* and *Junb*, which are involved in cytotoxicity and cytokine production. Lastly, Cluster 5 is strongly defined by the expression of the surface marker *Klra1*, which encodes the Ly49a receptor. Functional analysis supports marker gene’s functions as Cluster 3 is defined by response to stress and heat shock, Cluster 4 exhibits similar pathways to the non-responding Cluster 0 and Cluster 5 and is defined by cell-to-cell communication. Importantly, all three clusters -3, 4, and 5-display downregulation of interferon, interleukin, and cytokine signalling, also in line with a non-responding phenotype (**Supplementary** Figure 2C).

Surprisingly, none of the known NK cell receptors previously associated with memory NK cells to viruses (Azzi et al., 2014; Lopez-Vergès et al., 2011; Sun et al., 2009), such as NKG2D, Ly49D, Ly49H, Ly49F, or Ly49C, were specifically upregulated in any of the clusters of our dataset (**Supplementary** Figure 2E), in agreement with our previous observation (Camarasa et al., 2023), and suggesting a distinct mechanism for NK cell memory to *S. pneumoniae.* In addition, TLR receptors (*Tlr2* and *Tlr4)*, the main pattern recognition receptors for bacteria, are very lowly expressed, supporting our previous observation that they are not markers of the NK cMemory population to *S. pneumoniae* **(Supplementary** Figure 2D).

In summary, the analysis of NK cell clusters reveals distinct identities and functions. Cluster 0 indicates cells in steady state, while Cluster 1 is characterised by active immune engagement. Cluster 2, mainly composed of cMemory NK cells, shows readiness for future responses despite downregulated activity against current infections. Clusters 3, 4, and 5 represent non-responding phenotypes, emphasizing their different roles in immune regulation and communication.

### 3. Transcriptional reprogramming of cMemory NK Cells reveals distinct gene expression patterns following primary infection

The identification of Cluster 2, which is mainly populated by cMemory NK cells, shows that primary infection significantly modulates the transcriptional response of NK cells even when they are not actively responding to the infection. To further investigate whether cMemory NK cells are transcriptionally reprogrammed within individual clusters upon infection, we conducted differential expression analysis in the cMemory population compared to the cNaïve population in every cluster, with the aim to identify memory Markers (upregulated and downregulated genes) across clusters, and their distinct patterns of expression.

Differential expression analysis is graphically represented as upset plots representing upregulated or downregulated genes in cMemory cells for each cluster (**Figure 3A**). Horizontal bars (i.e., set size) correspond to the total number of differentially expressed genes per cluster, while vertical bars indicate the number of differentially expressed genes that are specific for a given cluster or shared across multiple clusters. The genes with highest fold changes are indicated above columns (**Figure 3**, **Tables 1-2, Supplementary table 1**). Several genes are uniquely upregulated or downregulated in a specific cluster, highlighting the different identity of cMemory NK cells in every cluster (**Figure 3A**, **Table 1**). Interestingly, the proportion of uniquely downregulated genes in specific clusters was higher in comparison to that of the upregulated genes (∼20% vs ∼10%), suggesting that the identity of cMemory cells in the different clusters is more defined by genes that are silenced than overexpressed. Notably, Cluster 1 is the most peculiar cluster. It has the lowest number of differentially expressed genes in its set size (23 downregulated and 14 upregulated) suggesting that there are less significant differences between cMemory and cNaïve NK cells within this cluster, probably indicating that transcriptional responses to infection are the dominant feature of this cluster.

**Figure 3.**
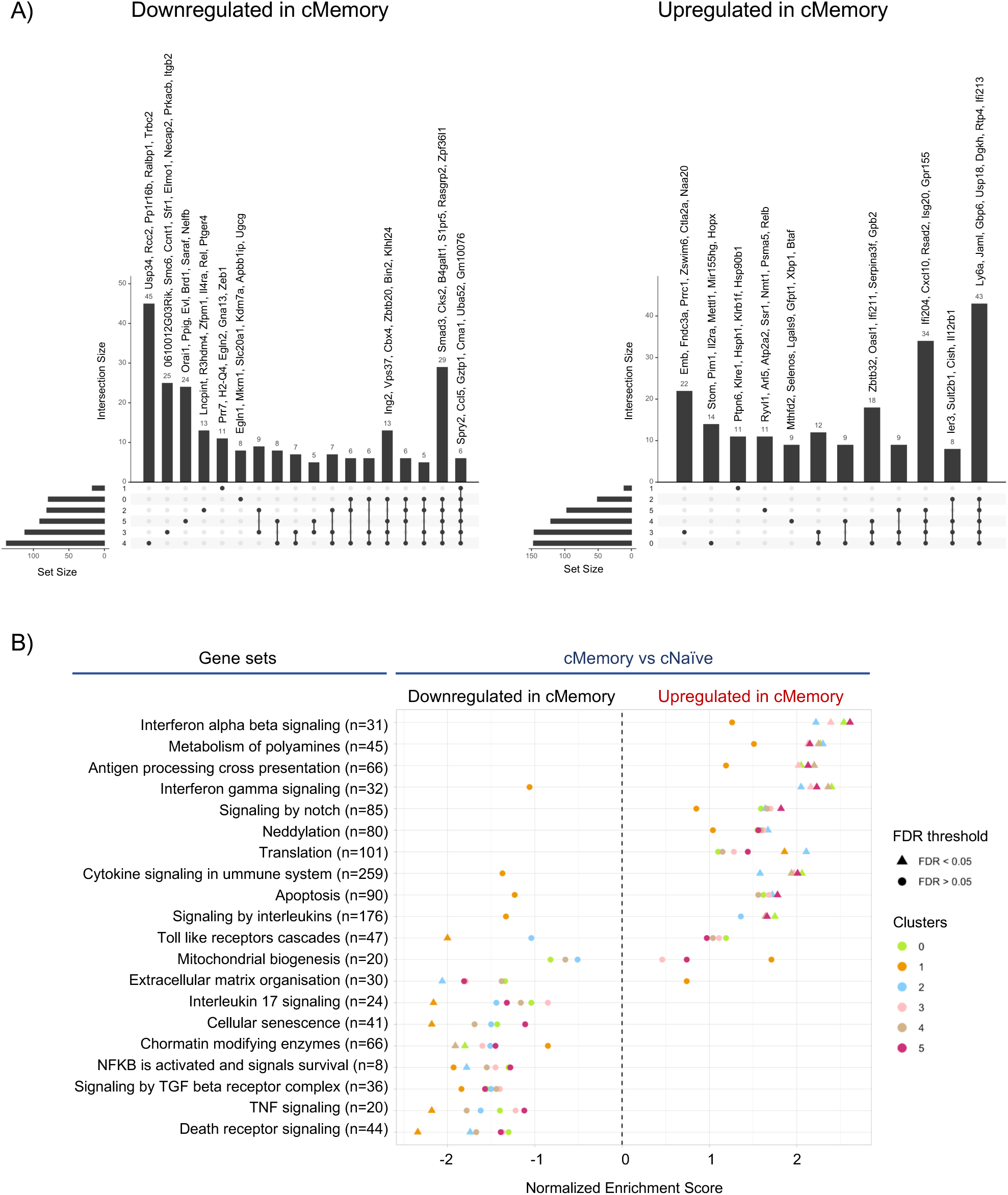
Transcriptional reprogramming of cMemory NK Cells reveals distinct gene expression patterns following primary infection. **A)** Upset plots representing differential expression analysis between cMemory and cNaïve cells. Horizontal bars represent the total number of significantly (adjusted p-value < 0.05 and absolute log fold change > 1.0) upregulated or downregulated genes per cluster. Vertical bars indicate the number of differentially expressed genes in one or more clusters. The genes with highest fold changes are indicated above columns. **B)** Gene Set Enrichment Analysis (GSEA) for upregulated or downregulated genes within the cMemory population in every cluster. Selected REACTOME genes set significantly enriched (adjusted p-value < 0.05) using the Normalized Enrichment Score (NES). N=Number of genes in every gene set.

**Table 1.** Differential state analysis: Genes upregulated in a specific cluster. List extracted from differential expression analysis between cMemory and cNaïve cells. Lists of genes are represented in the upset plots in Figure 3, in the columns where differentially expressed genes are specifically downregulated or upregulated in one cluster.

**Table 2.** Differential state analysis: Genes upregulated in an intersection of clusters. List extracted from differential expression analysis between cMemory and cNaïve cells. Lists of genes are represented in the upset plots in Figure 3, in the columns where differentially expressed genes are downregulated or upregulated in an intersection of clusters.

Importantly, we did not find any commonly upregulated genes between all clusters, and we found only 6 genes that are commonly downregulated (**Figure 3A**, **Table 2**). This suggests that there is a strong distinction between clusters, that there are no common markers expressed in all clusters that could define a memory signature, and that even the non-responding populations are reprogrammed during the primary infection. However, the intersection including clusters 0, 2, 3, 4 and 5 is the largest for both downregulated (29) or upregulated (43) genes suggesting that, excluding Cluster 1, the other clusters have certain common features. These include upregulation of activation markers (*Cd69, Ly6a, Ly6e, Xcl1*) or NK cell effector functions (*Ifng*), while the downregulated genes corresponded to cell migration markers (*S1pr5, Itgb7, Cd9, Adgre5*) (**Figure 3A**, **Table 2**).

This observation is aligned with the gene set enrichment analysis performed on the complete set of gene expression differences between cNaïve and cMemory NK cells within clusters (Figure 3B, Supplementary Table 2). Upregulation in cMemory NK cells is significantly enriched for interferon-alpha and beta signalling, metabolism of polyamines, antigen processing, and cross-presentation gene sets for all clusters except Cluster 1, which followed the same trend but not reaching significance. Interestingly, although the interferon gamma signalling gene set is upregulated in the cMemory population in all clusters but Cluster 1, expression of the Ifng receptor (*Ifngr1*) was downregulated (**Supplementary** Figure 3A). This aligns with our previous study showing that the protective memory phenotype is preserved in IfngrKO mice, suggesting that Ifng response is not the primary effector function of cMemory NK cells (Camarasa et al., 2023). For the interferon gamma signalling and other genes sets associated to the inflammatory response, such as cytokine signalling in immune system, apoptosis or signalling by interleukins, Cluster 1 follows an opposite trend with a downregulation for the cMemory NK cells. The reduced divergence between cMemory and cNaïve NK cells within Cluster 1 — identified in the cluster description analysis as a highly responsive cluster (Figure 2) — is likely due to both populations being actively engaged in the response to infection, thereby minimising functional differences. Other gene sets were enriched in downregulated genes across all clusters. These are related to cellular senescence, chromatin remodelling enzymes, and NF-κB-and TNF-signalling pathways (**Figure 3B**), suggesting an active inflammatory response associated with chromatin remodelling during primary infection.

In conclusion, the differential analysis of cMemory and cNaïve NK cells reveals that these populations not only have significant intrinsic transcriptional differences but also differ markedly in their upregulated and downregulated pathways, despite sharing similar cellular states within clusters. This supports the notion that following a primary infection, NK cells undergo transcriptional reprogramming, acquiring distinct identities across various cellular states.

### 4. Single cell trajectories reveal distinct transcriptional transitions between cMemory and cNaïve NK cells

To identify possible transitions between clusters driving the different responses in the cNaïve and cMemory populations, we performed cell trajectory analysis. In this analysis, a global lineage structure is estimated, which reflects the relative position of each cell within a biological transition, in this case, the NK cell response to *S. pneumoniae* infection. First, we conducted cell trajectory analysis in cNaïve (**Figure 4A, left panel**) and cMemory (**Figure 4A, right panel**) NK cells separately, defining Cluster 0 (the largest non-responding cluster) as the starting point of the trajectory. This analysis was complemented by examining the driving genes, whose expression levels change significantly along the trajectory axis or within specific branches. These genes are often key drivers of biological transitions and could shed light on which ones are most involved in orchestrating the NK cell response to *S. pneumoniae* infection.

**Figure 4.**
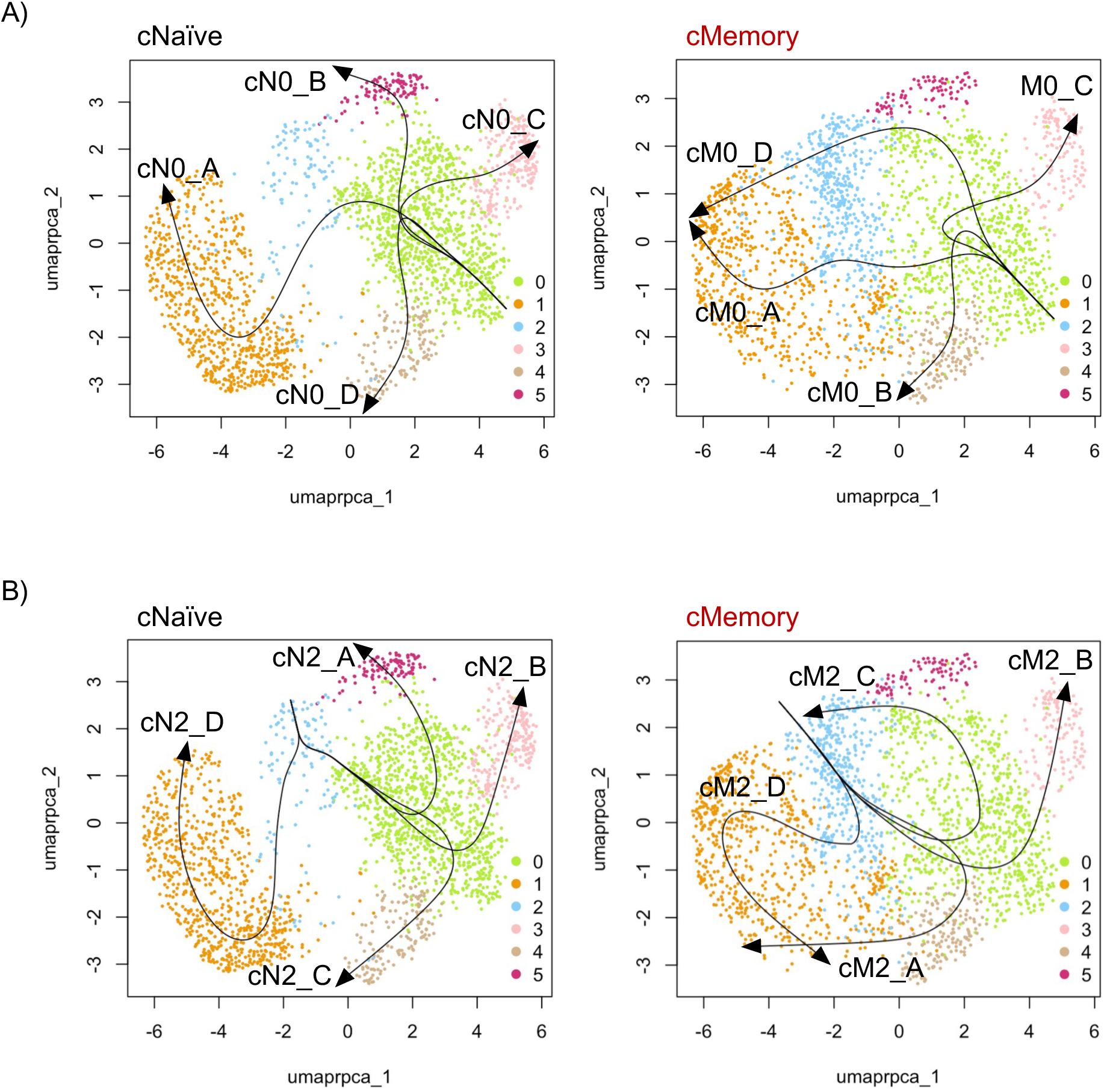
Single cell trajectories reveal distinct transcriptional transitions between cMemory and cNaïve NK cells A-B) Cell trajectory analysis in cNaïve (**left panel**) or cMemory (**right panel**) NK cells, defining Cluster 0 **(A)** or Cluster 2 **(B)** as the starting point of the trajectory.

4 distinct trajectories were identified (A through D) for both cNaïve and cMemory NK cells starting in cluster 0 (denoted as N0 or M0). All trajectories initially advance within Cluster 0 until they reach a branching point, where they diverge and acquire differential transcriptional programs. The N0_D and M0_B trajectories follow a similar path towards Cluster 4, while the N0_C and M0_C trajectories diverge slightly in their path towards Cluster 3. However, the N0_D/M0_B and N0_C/M0_C trajectories share only 9% and 4% of their top driving genes respectively (**Supplementary** Figure 4A), indicating that while the trajectories are similar, the genes driving them are distinct (**Table 3**). This further supports the finding that NK cells are reprogrammed following primary infection and highlights the intrinsic differences between cNaïve and cMemory NK cells within clusters.

**Table 3.** Top driving genes of single cell trajectories starting in Cluster 0. Top driving genes exclusively upregulated in every trajectory that was identified using cluster 0 as the starting point in the cMemory or cNaïve NK cells.

**Table 4.** Top driving genes of single cell trajectories starting in Cluster 1. Top driving genes exclusively upregulated in every trajectory that was identified using cluster 2 as the starting point in the cMemory or cNaïve NK cells.

Trajectories N0_A and M0_A share 14% of their top driving genes (**Supplementary** Figure 4A) being the most similar trajectories. These include the activation marker *Ly6a*, the transcription factors *Nr4a2, Ier2*, and *Atf6*, as well as immune response signalling genes such as *Spry2, Adgre5*, and *CD53*. However, the proportion of uniquely expressed genes is 26% and 27% respectively (**Supplementary** Figure 4A), suggesting that they are more diverse than similar. Functionally, the top driving genes in trajectory N0_A are involved in immune response, cell-adhesion, and apoptosis, while top driving genes of the M0_A trajectory are related to immune response and lymphocyte differentiation (**Table 3, Supplementary** Figure 4C).

Interestingly, N0_B and M0_D follow very distinct trajectories: both progress towards Cluster 5 but N0_B stops there, while M0_D continues towards Cluster 2 and subsequently Cluster 1. Additionally, M0_D shares only one top driving gene with N0_A (*Zeb2*), indicating that M0_D represents a unique trajectory that emerges specifically in cMemory NK cells, encompassing Cluster 2 (**Figure 4A, right**). In addition, this trajectory joins M0_A that links Cluster 0 (non-responding) and Cluster 1 (responding) running through Cluster 2, which is composed of 80% cMemory NK cells and could represents an intermediary cell population. Strikingly, trajectories M0_D and M0_A describe a circular transition that could be happening in a recurrent manner between non-responding and responding NK cMemory cell states. The top driving genes of the M0_D trajectory (**Supplementary** Figure 4C **and Table 3**) are involved in rRNA processing, ribosome biogenesis, mitochondrial function **(Supplementary** Figure 4C), and include genes linked to immune regulation, suggesting a population of dividing and metabolically active cells, with a subset likely in a preparatory phase for cell division.

To better understand the origin and fate of cells in Cluster 2, we performed a similar trajectory analysis but now defining Cluster 2 as the starting point (**Figure 4B**). Again, 4 distinct new trajectories were identified (A through D) for both cNaïve and cMemory NK cells (denoted as N2 or M2). Trajectories in the cNaïve population started in a less dense Cluster 2 (20% of the whole Cluster) and diverged towards the other clusters. Trajectories N2_A, N2_B and N2_C passed through Cluster 1 before moving towards clusters, 5, 3 and 4 respectively. Trajectory N2_D went directly from Cluster 2 to Cluster 1, suggesting a population of cells transitioning towards active immune response. Indeed, functional analysis of trajectory N2_D shows an enrichment for pathways linked to response to infection and apoptosis **(Supplementary** Figure 4C).

Importantly, trajectories in the cMemory population were very different. Only trajectory M2_B was similar to N2_B even though, again, only 5% of top driving genes were common between the two trajectories. Interestingly, trajectories M2_D and M2_A also showed a circular pattern, probably representing a recurring cell state transition through clusters 2, 0, 4 and 1. This suggests a continuity between the cMemory-enriched Cluster 2, the non-responsive Cluster 0 and the responsive Cluster 1, or in the other sense. In addition, trajectory M2_C forms a continuous closed-loop structure, also indicating a circular progression between Cluster 2 and Cluster 1 and suggesting that cells in Cluster 2 could arise from Cluster 0, or vice versa. Functional analysis of the M2_D trajectory includes response to viruses, negative regulation of defense responses or apoptosis while active immune response, rRNA processing and viral response is enriched in the M2_A trajectory. Both M2D and M2A transition towards a functionally active response. However, functional analysis for M2_C trajectory shows enrichment for pathways related to ribosome biogenesis, but not to an active response to infection. This in line with an oscillation between a steady-state and an active state between Cluster 0 and Cluster 1, with Cluster 2 as an intermediate state for cMemory NK cells.

Notably, the phenomenon of circularity of the trajectories is only found in the cMemory population, which could align with the fact that cMemory NK cells have already changed their basal state following the first infection (Cluster 2) and that cells in Cluster 2 either alternate with cells in Cluster 0 (which are non-responding but more active that the cNaïve ones) or with cells in Cluster 1, a more general responsive state.

### 5. cMemory NK cells display enhanced activation and proliferation capacity following *S. pneumoniae* infection

Reported properties of cMemory NK cells are increased activation markers (Romee et al., 2012; Wijaya et al., 2021) and cell division (Adams et al., 2019; Grassmann et al., 2019). Therefore, we first analysed the expression of specific markers of NK cell activation in our scRNA-seq dataset and represented individual gene expression throughout the clusters (**Figure 5A**). Expression of *Cd69*, a gene encoding a classical surface marker of NK cell activation, is significantly elevated in cMemory NK cells compared to cNaïve NK cells across all clusters and in the global analysis of all cMemory versus cNaïve NK cells (**Figure 5B, top; Supplementary** Figure 5A). *Ly6a* (also known as *Sca-1*), encoding for a surface protein specifically expressed on activated lymphocytes (Stanford et al., 1997) and an early marker of NK cell activation (Fogel et al., 2013), is also significantly upregulated in the cMemory NK cell population consistently across all individual clusters, and in the global analysis (**Figure 5B, mid panel; Supplementary** Figure 5B**, top**). Similarly, *Ly6e*, which encodes another surface protein associated cell activation (AlHossiny et al., 2016), is consistently upregulated in cMemory NK cells (**Figure 5B, middle panel; Supplementary** Figure 5B**, bottom**). Notably, the combined upregulation of *Cd69, Ly6a, Ly6e*, is highly increased in cMemory NK cells (**Supplementary** Figure 5C), reinforcing their activated state and potential effector capacity.

**Figure 5.**
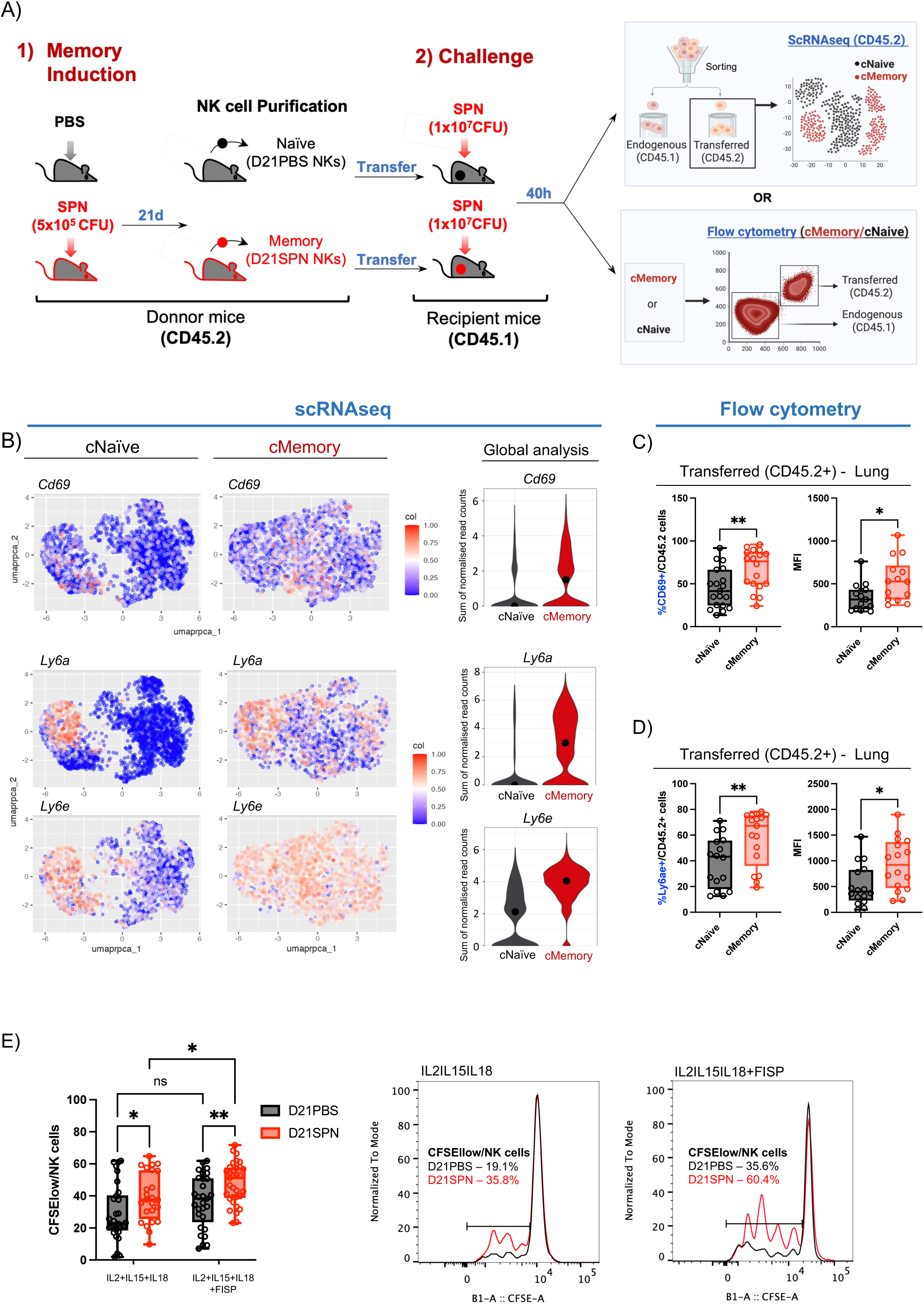
Enhanced activation and proliferation capacity of cMemory NK cells in response to *S. pneumoniae* infection. **A)** Experimental scheme for model set up as described in Figure 5A for scRANseq. Alternatively, 40h aftertransfer and challenge of the recipient mice, cell suspensions of bulk NK cells from recipient mice (including endogenous CD45.1 and transferred CD45.2 NK cells) were prepared for ex vivo staining and FACS analysis. **B)** Panel plot representing cNaïve or cMemory NK cells in an UMAP plot representation where cells are coloured by *Cd69, Ly6a* or *Ly6e* expression (**left panel**). Violin plots representing the sum of the normalised read counts for *Cd69, Ly6a* or *Ly6e*, in the total of cNaïve or cMemory NK cells. **C, D, E)** C57BL/6 mice were intranasally infected with either PBS (black symbols) or sub-lethal dose of S. *pneumoniae* After 21 days, NK cells were highly purified from spleens of D21PBS or D21SPN mice and transferred to recipient mice **(C and D)** or cultured *in vitro* **(E)**. **C, D)** Flow cytometry analysis of the percentage of CD69+ NK cells **(C)** or Ly6ae+ NK cells **(D)** among transferred (CD45.2+) cNaïve or cMemory NK cells and MFI. Each dot in the boxplots represents an individual mouse (black dots for mice having received D21PBS NKs and red dots for mice having received D21SPN NKs), lines are the median and error bar show min to max. Data are pooled from ≥ 3 repeats with n≥3 mice/group. **E)** D21PBS and D21SPN purified NK cells were pooled from n≥3 mice, stained with CFSE and incubated *in vitro* in n≥3replicates/group with cytokines (IL-15 at 2 ng/ml, IL-18 at 1,5 ng/ml, IL-2 at 1000U/ml) for 4 days, with or without formaldehyde-inactivated bacteria, at a bacteria-to-cell ratio of 40:1 during the last 24 hours. Flow cytometry analysis of the percentage of CFSE^low^ NK cells among cultured D21PBS or D21SPN NK cells. Boxplot where each dot represents an experimental replicate (black dots for D21PBS NK cells, red dots for D21SPN NKs cells), lines are median, error bar show min to max. Data are representative n≥3 experiments (**left panel**). Representative overlay histogram of D21PBS (black) and D21SPN (red) NK cells upon IL2-IL15+IL18+SPN stimulation without (**middle panel**) or with fixed *S. pneumoniae* (FISP) stimulation (**right panel**). ns, not significant. * p < 0.05 and ** p <0.01, t-test with crossed design for single comparisons and 2way ANOVA test for multiple comparisons.

We also analysed specific genes showing significant upregulation *in vivo* (**Figure 5A**). Following the same experimental setting, lungs and spleens were harvested from recipient mice 40h post challenge. Then, cell suspensions of bulk NK cells from recipient mice (including endogenous CD45.1 and transferred CD45.2 NK cells) were prepared for ex vivo staining and FACS analysis. The proportion of CD69+ cells is significantly increased in transferred (CD45.2⁺) cMemory NK cells in the lung, along with an increase in their mean fluorescence intensity (MFI) (**Figure 5C**). To assess Ly6a and Ly6e levels, we used a common antibody recognising both proteins and observed Ly6ae+ cells and their MFI are also significantly increased in these cMemory NK cells (**Figure 5D**). Therefore, both at the transcriptional and protein level, cMemory NK cells display higher levels of activation markers in lung. To understand if this is a local or systemic effect we also analysed transferred (CD45.2+) NK cells in the spleen and, interestingly, the percentage of CD69+ or Ly6ae+ cells is generally higher for both cNaïve and cMemory NK cells, however, a similar trend of increased percentage and MFI is observed in cMemory NK cells (**Supplementary** Figure 5D). Notably, endogenous NK cells (CD45.1⁺) also show increased frequencies and MFI of CD69+ and Ly6a/e+ in both the lung and spleen in mice having received Memory NK cells (**Supplementary** Figure 5E). Surprisingly, these results suggest that transferred cMemory cells confer their activated status to surrounding endogenous NK cells. Collectively these findings suggest that the presence of cMemory NK cells facilitates a faster and more robust activation of both transferred and endogenous NK cells during *S. pneumoniae* infection.

Next, as Memory NK cells have been shown to have a high expansion capacity, D21PBS and D21SPN NK cells were stained with CFSE to monitor cell division and incubated *in vitro* for 4 days with IL-15 and IL-18 to sustain NK cell viability and IL-2 to promote proliferation. Under these conditions, D21SPN (Memory) NK cells display a higher percentage of cells having diluted CFSE through cell division (CFSE^low^) compared to D21PBS (Naïve) NK cells. Importantly, this difference is significantly enhanced when cells are challenged with PFA fixed *S. pneumoniae* (FISP) (**Figure 5E**) suggesting an increased proliferative capacity of D21SPN (Memory) NK cells in response to the infection. *In vivo*, the absolute number of NK cells following challenge in mice having received Memory NK cells is significantly increased in the lung and spleen compared to mice having received Naïve NK cells (Supplementary Figure 5F), supporting the increased proliferation capacity of cMemory NK cells observed *in vitro*. In addition, we performed Ki67 staining and observed an increased percentage of Ki67+ NK cells in the mice having received Memory NK cells (Supplementary Figure 5G) in the lung. Interestingly, we observed the opposite effect in the spleen suggesting that cMemory NK cells in the spleen might have stopped proliferating 40h post-challenge or have migrated to other organs.

### 6. cMemory NK cells exhibit enhanced residency and survival phenotype compared to cNaïve NK cells

One of the main characteristics of NK cells is their circulating capacity, as they can be recruited to organs to safeguard tissue functions. We thus analysed the expression of genes coding for circulation (*Klr2, Ifngr1, Sell, Zeb2, S1pr5, Cxcr3, Klrg1, Hhex, Hes1, Itga2*) or residency markers (*Itga1, Cd69, Rgs2, Vps37b, Rgs1, Tnfaip3, Bhlhe40, Xcl1, Nffibia, Pim1*) in NK cells as previously reported (Crinier et al., 2018; Torcellan et al., 2024). Interestingly, circulating markers are depleted in Cluster 1 in both cNaïve and cMemory populations, (**Figures 6A**) suggesting that NK cells responding to infection have a more tissue resident phenotype. In addition, in the other clusters, genes linked to NK cell residency phenotypes are upregulated while circulating genes are downregulated, suggesting that, in general, cMemory NK cells are more resident than circulating.

**Figure 6.**
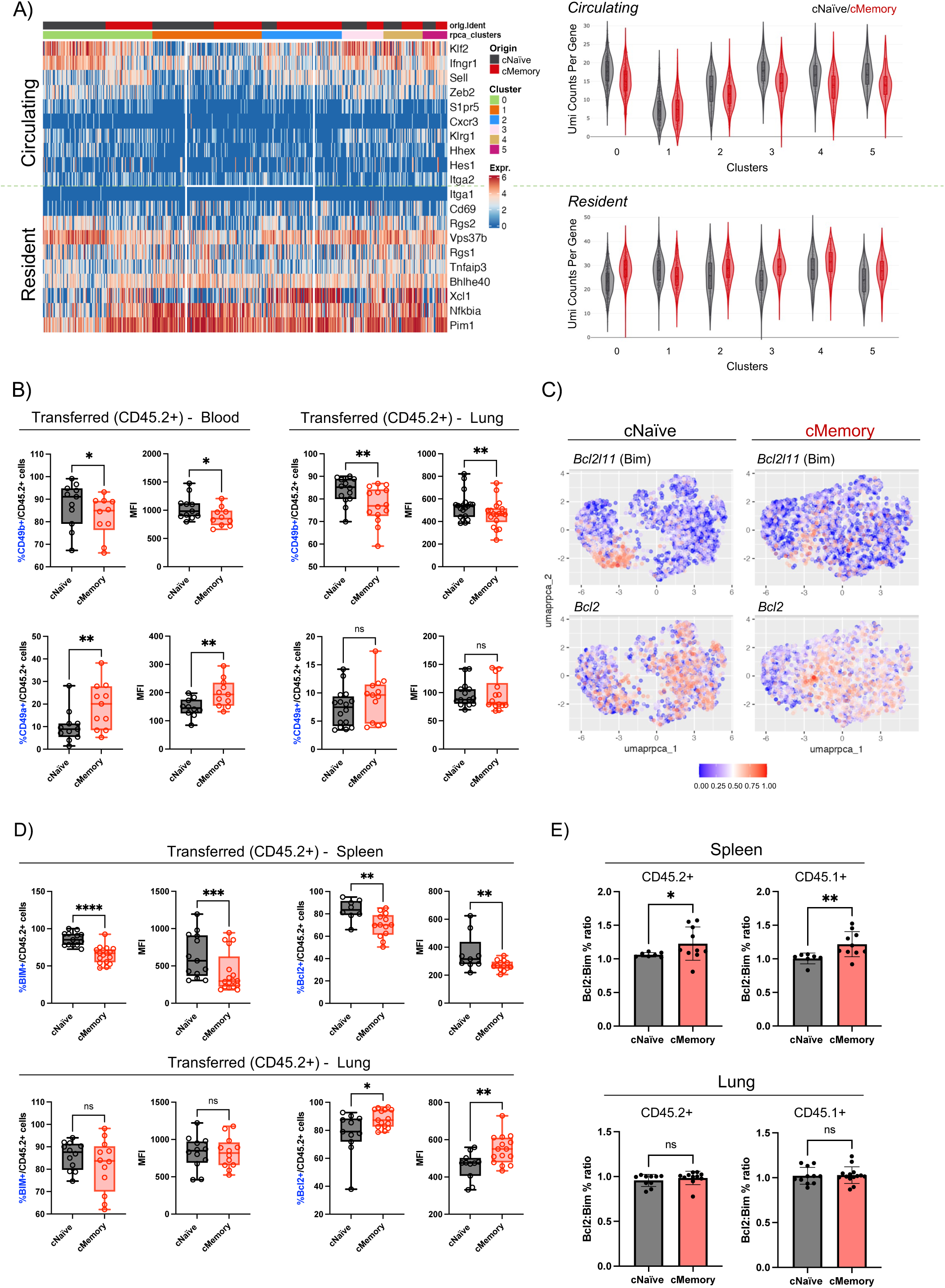
cMemory NK cells exhibit enhanced residency and survival phenotype. **A)** Heatmap representing gene counts for the expression of circulating or resident markers within clusters, divided by cNaïve or cMemory NK cells (**left panel**). Violin plots representing the sum of the normalize gene counts per circulating or resident markers in the cMemory or cNaïve population within every cluster (**right panel**). **B,D,E)** C57BL/6 mice were intranasally infected with either PBS (black symbols) or sub-lethal dose of S. After 21 days, NK cells were highly purified from spleens of D21PBS or D21SPN mice and transferred to recipient mice. **B)** Flow cytometry analysis of the percentage of Cd49b+ NK cells **(top panel)** or Cd49a+ NK cells **(low panel)** among transferred (CD45.2+) cNaïve or cMemory NK cells in blood or lung and its MFI for DX5 or CD49a. Each dot in the boxplots represents an individual mouse (black dots for mice having received D21PBS NKs and red dots for mice having received D21SPN NKs), lines are the median and error bar show min to max. **C)** Panel plot representing cNaïve or cMemory NK cells in an UMAP plot representation where cells are coloured by *Bcl2l11* (Bim) (top) or *Bcl2* expression (bottom). **D)** Flow cytometry analysis of the percentage of Bim+ NK cells or Bcl2+ NK cells among transferred (CD45.2+) cNaïve or cMemory NK cells in spleen **(top)** or lung **(bottom)** and its MFI for Bim or Bcl2. **E)** Ratio of Bcl2:Bim positive cells among transferred (CD45.2+) cNaïve or cMemory NK cells or among endogenous (CD45.1+) NK cells in spleen (**top**) or lung (**bottom**) of mice having received cNaïve or cMemory NK cells. Each dot in the boxplots represents an individual mouse (black dots for mice having received D21PBS NKs and red dots for mice having received D21SPN NKs), lines are the median and error bar show min to max. Data are pooled from ≥ 3 repeats with n≥3 mice/group. ns, not significant. * p < 0.05 and ** p <0.01, t-test with crossed design.

To confirm these results *in vivo*, we used two well-described surface markers for circulation and residency in NK cells: Cd49b (DX5) as a circulation marker and Cd49a as a residency marker. The results confirm that transferred cMemory NK cells (CD45.2+) display a more resident phenotype, as evidenced by a significant decrease in the percentage of Cd49b+ cells in both the blood and lung (**Figure 6B**). In addition, the percentage of CD49a+ cells is increased in blood, also indicating a more resident phenotype (**Figure 6B**). A similar but not significant trend was observed in endogenous (CD45.1+) NK cells in blood and lung (**Supplementary** Figure 6A) and in transferred and endogenous cells in spleen (**Supplementary** Figure 6B).

NK cell persistence in tissues is tightly linked to survival mechanisms. In NK cells, apoptosis and survival are regulated by the balance between the anti-apoptotic factor Bcl-2 and the pro-apoptotic factor Bim (Min-Oo et al., 2014; Riggan et al., 2022). In our analysis*, Bcl2l11* (coding for Bim) expression is globally downregulated in cMemory NK cells (**Figure 6C, Supplementary** Figure 6C) but is significantly upregulated in cNaïve NK cells within Cluster 1 (**Supplementary** Figure 6D). This suggests an impaired upregulation of pro-apoptotic signalling in cMemory NK cells, particularly within Cluster 1 during active responses. Conversely, *Bcl2* expression is globally upregulated in cMemory NK cells (**Figure 6C, Supplementary** Figure 6C) and consistently elevated across all clusters (**Supplementary** Figure 6D). Interestingly, no significant differences in Bim levels are observed *in vivo* in transferred (CD45.2⁺) cells within the lung, however, we detect a significant reduction in both the number of Bim+ cells and MFI in the spleen (**Figure 6D**). In contrast, Bcl2 is downregulated in the spleen, but it is significantly increased in the lung (**Figure 6D**). This imbalance resulted in a significantly increased Bcl-2:Bim ratio in the spleen but not in the lung (**Figure 6E**), indicating enhanced anti-apoptotic signalling via Bcl-2 in the spleens of mice that received cMemory NK cells. The enhanced survival and residency features of cMemory NK cells could contribute to their long-term presence in tissues, allowing them to provide a rapid, sustained immune defence against re-infection.

### 7. cMemory NK cells have intrinsically increased cytotoxic activity

Having observed in a previous study the potential importance of cytotoxic activity in the response of Memory NK cells, we next focused on the analysis of Granzyme B (*Gzmb*) and Perforin (*Prf1*) expression across distinct NK cell clusters. *Gzmb* expression is increased in the responding Cluster 1 for both cNaïve and cMemory NK cells, with a significant increase in cNaïve NK cells. Conversely, in other clusters, *Gzmb* expression is significantly higher in cMemory NK cells, suggesting a general upregulation in Memory NK cells across clusters, but with a specific elevation in cNaïve NK cells within the responding Cluster 1, corresponding to cells actively responding to the infection for the first time (**Figure 7A, top**). In contrast, *Prf1* expression is uniformly upregulated in cMemory NK cells in general and across all clusters **(Figure 7A, bottom**). Interestingly, the expression of Granzyme B and Perforin appears to be uncoordinated among NK cells across the clusters, highlighting potential differences in their regulation. In addition, our analysis reveals that while there is an overall increase in cMemory NK cells co-expressing both *Gzmb* and *Prf1* (**Figure 7B, top panel**), Cluster 1 displays a higher proportion of cells expressing either *Gzmb or Prf1* individually while other clusters exhibit an increased co-expression of both molecules (**Figure 7B, bottom panel**).

**Figure 7.**
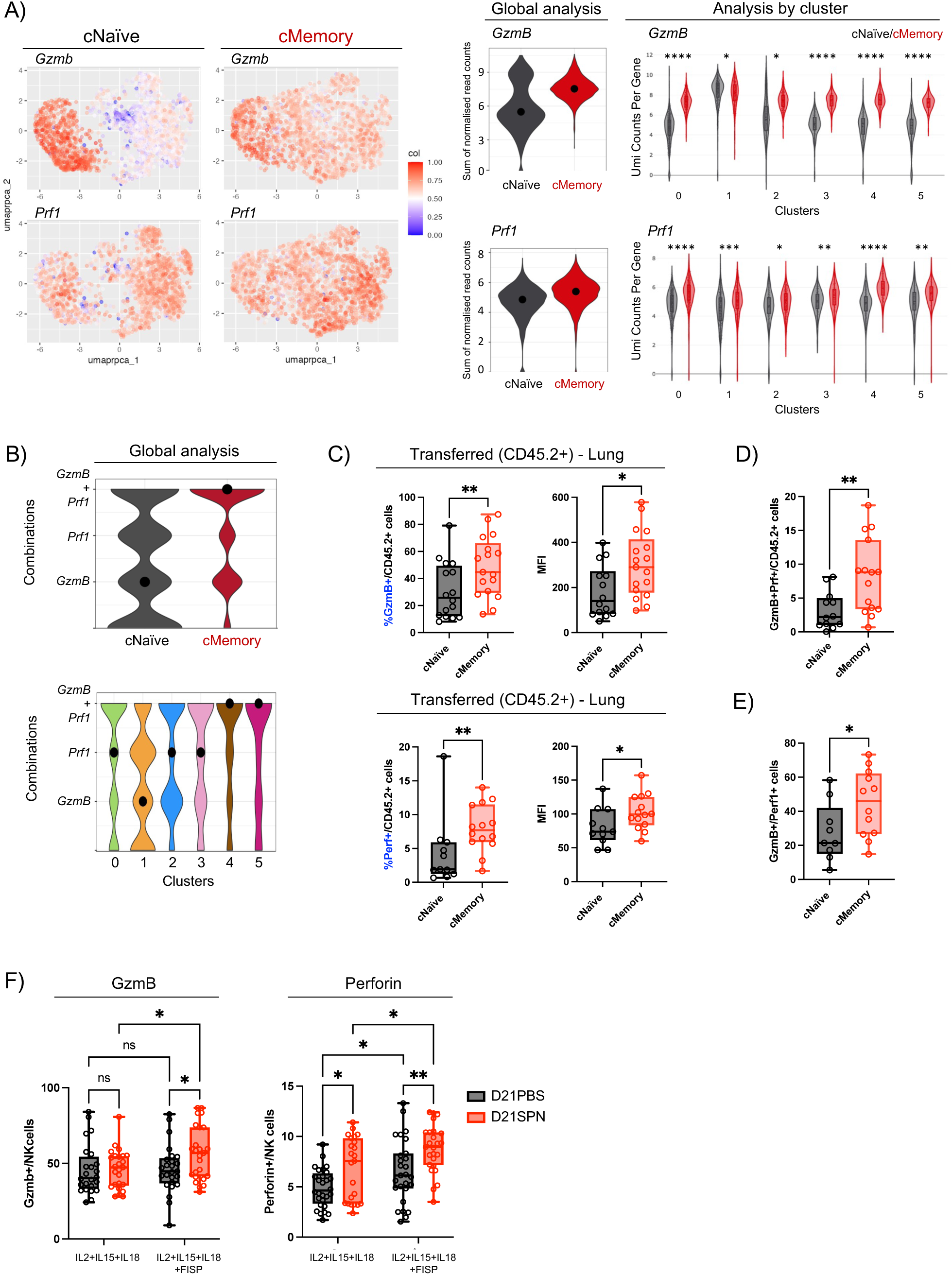
cMemory NK cells have intrinsically increased cytotoxic activity. **A)** Panel plot representing cNaïve or cMemory NK cells in an UMAP plot representation where cells are coloured by *Gzmb* or *Prf1* expression **(left panel)** and violin plots representing the sum of the normalised read counts for in the total of cNaïve or cMemory NK cells **(mid panel)** or within every cluster **(right panel)**. **B)** Combinations of normalized counts for *GzmB and Perforin* in cNaïve or cMemory NK cells **(top panel)** and in the different clusters (bottom panel). **C,D,E)** C57BL/6 mice were intranasally infected with either PBS (black symbols) or sub-lethal dose of S. After 21 days, NK cells were highly purified from spleens of D21PBS or D21SPN mice and transferred to recipient mice (**C,D, E**) or cultured *in vitro* (**F**). Flow cytometry analysis of the percentage of GzmB+ NK cells **(B)** or Perf+ NK cells **(C)** among transferred (CD45.2+) cNaïve or cMemory NK cells in lung and their MFI. **F)** D21PBS and D21SPN purified NK cells were pooled from n≥3 mice and incubated *in vitro* in n≥3 replicates/group with cytokines (IL-15 at 2 ng/ml, IL-18 at 1,5 ng/ml, IL-2 at 1000U/ml) for 4 days, with or without formaldehyde-inactivated bacteria, at a bacteria-to-cell ratio of 40:1 during the last 24 hours. Flow cytometry analysis of the percentage of Gzmb+ **(left)** or Perf+ (**right**) NK cells among cultured D21PBS or D21SPN NK cells. Boxplot where each dot represents an experimental replicate (black dots for D21PBS NK cells, red dots for D21SPN NKs cells), lines are median, error bar show min to max. Data are representative n≥3 experiments (**left panel**). ns, not significant. * p < 0.05 and ** p <0.01, t-test for crossed design for single comparisons and 2way ANOVA test for multiple comparisons.

Given the significant upregulation of *Gzmb* and *Perf1* genes in cMemory NK cells, we investigated whether their expression levels correlated with global protein levels *in vivo*. To address this, we analysed transferred (CD45.2+) challenged cNaïve and cMemory NK cells by flow cytometry. The results reveal a significant increase in the percentage of Granzyme B+ and Perforin+ cMemory NK cells, along with a marked increase in Granzyme B mean fluorescence intensity (MFI) (**Figure 7C**). Interestingly, the *in vivo* percentages of Granzyme B+ and Perforin+ cells did not correlate with each other, reinforcing the notion of their uncoordinated expression. These findings align with the scRNA-seq results, as recipient mice with transferred Memory NK cells show an increase in double-positive cells; however, these account for only ∼10% of the total NK cell population (**Figure 7D**). Notably, within the Perforin+ subset, approximately 50% also express Granzyme B (**Figure 7E**). Additionally, we analysed transferred NK cells in the spleen (**Supplementary** Figure 7A) and observed a similar increase in both Perforin in Memory NK cells, consistent with a systemic infection. Finally, we assessed Granzyme B and Perforin levels in endogenous (CD45.1+) NK cells within the lung and spleen (**Supplementary** Figure 7B). Surprisingly, endogenous NK cells follow similar trends as the transferred cells, with increased expression of both cytotoxic molecules. These findings suggest that transferred Memory NK cells are sufficient to enhance the cytotoxic response of endogenous NK cells, highlighting a systemic effect on the endogenous host NK cell population.

To determine whether the increased expression and production of cytotoxic proteins was an intrinsic property of NK cells, we performed *in vitro* studies using purified D21PBS and D21SPN cells challenged with FISP. NK cells were maintained for 4 days in the presence of IL-15, IL-18, and IL-2, followed by a 24-hour exposure to FISP. In the absence of SPN, D21PBS (Naïve) and D21SPN (Memory) NK cells present comparable percentages of Granzyme B+ cells. However, FISP challenge results in a significant increase in the percentage of Granzyme B+ cells (**Figure 7F, left**). In contrast, Perforin+ D1SPN NK cells increase both before and after FISP challenge, and significantly increase in response to FISP. These results indicate that while Perforin expression is constitutively upregulated in D21SPN (Memory) NK cells, it can be further enhanced by *S. pneumoniae* stimulation. Together, these findings highlight the uncoordinated regulation of Granzyme B and Perforin as an intrinsic feature of Memory NK cells.

Finally, we tested whether NK cell degranulation could kill or restrict *S. pneumoniae growth*. We performed *in vitro* experiments where we incubated live *S. pneumoniae* with the supernatants Naïve or Memory NK cells incubated with cytokines and with or without FISP *in vitro* (**Supplementary** Figure 7C). After 4 hours of incubation, we did not observe a difference in bacterial counts in supernatants from Naïve or Memory NK cells that were not stimulated with FISP. However, we observed a small decrease of bacterial counts when these were incubated with supernatants from Memory cells incubated with FISP. These results are in line with the significant increase of GzmB+ NK cells following FISP stimulation and suggest that NK cell degranulation can restrict *S. pneumoniae* growth.

## DISCUSSION

The potential of using NK cell memory to combat infections holds promise, however, cellular responses and memory functions to bacterial infections are only starting to emerge. Although functionality of memory NK cells against *S. pneumoniae* has been exposed, here, we characterize the emerging Memory NK cell populations and their respective transcriptional programs. Our analysis firmly establishes the appearance of functionally diverse NK cell subpopulations following infection with *S. pneumoniae* **(Supplemetary Figure 7D).**

Differential analysis and lineage tracing of cNaïve and cMemory NK cells reveal not only significant intrinsic transcriptional differences between cNaïve and cMemory NK cells across all clusters, but also that despite sharing similar cellular states within clusters, cNaïve and cMemory NK cells differ markedly in their upregulated genes. However, both cMemory and cNaïve cells within Cluster 1 present a stronger response in many aspects (decreased Ifng or Gzmb expression, less circulation markers,…) than cMemory cells in other clusters. In fact, it is possible that not all cMemory NK cells have acquired a protective phenotype during the primary infection and are not fully sensitised to the infection, providing a primary-like response during the challenge (**Supplementary** Figure 7D). Still, genes like Gzmb or Ifngr1 are significantly downregulated in the cMemory NK cells in Cluster 1 compared to the cNaïve NK cells in Cluster 1, suggesting a more moderate response of these cells in response to the secondary infection. These results support that NK cells, following a primary infection, undergo transcriptional reprogramming acquiring distinct identities across various cellular states, and providing different response profiles upon re-infection. Further supporting this hypothesis, lineage tracing of cMemory NK cells reveals recurrency trends between the highly responding (Cluster 1) and less-responding states (Cluster 0), with an intermediary Memory state (Cluster 2). Importantly, this pattern is exclusively found in the cMemory population, in correlation with a primary reprogramming of these cells and a reversible cellular state as a property of their memory identity. The roles of these diverse cMemory subpopulations remains to be investigated, with the possibility that a cMemory population remains unresponsive (Cluster 2) to protect its longed-lived phenotype, while other populations provide a more effective and protective immune response, as it has been suggested in other models (Adams et al., 2019; Schuster et al., 2023a).

Memory phenotypes in innate immune cells have been correlated with the acquisition of permanent chromatin rearrangements, that provide maintenance to the memory identity and function. NK cell memory subpopulations have also been described to correlate with chromatin rearrangements in various models (Lau et al., 2022). We have previously shown that NK cell memory to LPS or to *S. pneumoniae* correlates with the enrichment of H3K4me1 at enhancers of *Gzmb* and *Ifng* (Camarasa et al., 2023; Rasid et al., 2019). In the present study, we have identified different transcription factors as marker genes of Cluster 1 (highly responding) or Cluster 2 (enriched in cMemory NK cells) (**Figure 2B**) suggesting different regulation of their transcriptional activity at the level of chromatin. In addition, differential analysis revealed that these were differently expressed in the cNaïve or cMemory population within these clusters (Table 1, data not shown), suggesting that they might play a specific role in the cMemory or in the cNaïve NK cells. Potentially, some transcription factors could be contributing to the Memory establishment during the primary response in the Naïve NK cells, while others could be more involved in its maintenance within the Memory population. This hypothesis aligns with previously described memory mechanisms that speculate that the maintenance of the accessibility of memory domains might be driven by secondary TFs that access these domains following their opening by pioneering TFs during a first stimulation (de Laval et al., 2020; Larsen et al., 2021; Lau et al., 2018; Naik et al., 2017; Ordovas-Montanes et al., 2018). Indeed, one of the TFs that is significantly upregulated in cMemory NK cells across most clusters is Atf3, which has been described to remain bound to memory domains following the opening of these domains after a first stimulation (Larsen et al., 2021). Further studies will be needed to understand the specific mechanisms of acquisition and maintenance of NK cell memory to *S. pneumoniae* at the transcriptional level and at the level of chromatin, and how long-term changes can drive a differential response in the cMemory NK cells.

Numerous studies have reported the existence of tissue-resident NK cells subpopulations in mice and humans (Crinier et al., 2018; Sojka et al., 2014; Sparano et al., 2024; Tessmer et al., 2011) but also the appearance of new memory subpopulations that establish residency in different organs upon viral infection (Flommersfeld et al., 2021; Schuster et al., 2023b; Torcellan et al., 2024). This supports the idea that NK cells are not only circulating lymphocytes that infiltrate infected tissues, but that memory subpopulations of tissue-resident NK cells are permanently established in specific tissues for an enhanced protective action. Our results are in agreement with this idea and show that responding cMemory NK cells present a more tissue-resident phenotype. In addition, pro-survival genes are upregulated across most clusters (*Bcl2, Gimap7, Socs1*), and we have observed an anti-apoptotic phenotype in cMemory NK cells in the spleen, suggesting that the combination of enhanced survival and residency features of cMemory NK cells could contribute to their long-term presence in tissues, allowing them to provide a rapid, sustained immune defence against re-infection.

Interestingly, our results show that upon transfer of Memory NK cells and challenge of the recipient mice, endogenous NK cells respond in a similar manner, suggesting that transferred cMemory NK cells might stimulate endogenous NK cells to ensure a coordinated response. While some studies have shown how trained innate immune cells can influence the activity of other immune cells to produce a more generalised and enhanced response (Kleinnijenhuis et al., 2012; Quintin et al., 2012; Saeed et al., 2014) direct evidence of them inducing a trained or memory phenotype in naïve cells of the same type is currently lacking. Although in our previous study(Camarasa et al., 2023) we did not observe a significant increase in other immune cell populations in challenged mice receiving Memory NK cells, the transferred cMemory NK cells exhibited elevated *Ifng* expression, which could stimulate cytokine production by other immune cells and promote the activation of endogenous NK cells. Additionally, the overexpression of chemokines such as *Cxcl1*0 may contribute to the recruitment of NK cells to sites of infection, enhancing their exposure to activating cytokines. In addition, it remains to be investigated if the stimulation of endogenous NK cells is induced during the response, or if transferred NK cells can influence the phenotype of endogenous NK cell following their transfer into the recipient mice. Further research will be needed to explore the potential for such intercellular training mechanisms in NK cells.

Our transcriptional data reported here shows that cMemory NK cells globally upregulate both Granzyme B and Perforin, providing a stronger cytotoxic response against reinfection with *S. pneumoniae*. We have confirmed that an increase in Granzyme B and Perforin is also quantifiable at the protein level during infection *in vivo*. A similar induction is induced *in vitro* by the addition of fixed *S. pneumoniae*, which demonstrates a NK cell intrinsic response to direct sensing of *S. pneumoniae*. In addition, supernatants of stimulated NK cells decreased live *S. pneumoniae* bacterial counts. These results suggest a direct activity of either or both Granzyme B and Perforin on bacteria, however neither bacterial recognition nor direct killing has ever been demonstrated. Other reports showed a reduction of bacterial kinetics when bacteria, such as *E. coli* and *B. cenocepacia*, were incubated *in vitro* with human NK cells, but the mechanism at play is unknown (Garcia-Peñarrubia et al., 1989; S. S. Li et al., 2019).

Our *in vitro* experiments demonstrate that the increased cytotoxicity of responding Memory NK cells is an intrinsic property encoded in their transcriptional program. However, how NK cells manage to sense *S. pneumoniae* and provide a response remains to be elucidated. In our scRNA-seq analysis, we did not find increased expression of genes coding for NK cell activating receptors involved in viral recognition such as Ly49 and NKG2 families or well-known pattern recognition receptors such as TLR2 or 4. These results are in agreement with our previous work (Camarasa et al., 2023; Rasid et al., 2019). Therefore, we can hypothesise that another type of cell surface protein is involved in recognising pneumococci. Interestingly, we found a general upregulation of the surface markers Ly6a and Ly6e in cMemory NK cells. Ly6A (also called Sca-1) is found in diverse murine cell types including hematopoietic stem cells and lymphocytes (Upadhyay, 2019). One study showed that Ly6A was highly upregulated during MCMV infection, and the authors suggested that it can serve as a marker of early and nonselective NK cell activation (Fogel et al., 2013). Ly6E plays a redundant role in thymic development and development of immune cells. However, expression and function of these receptors on NK cells are poorly understood. Another surface marker that is upregulated in the cMemory population is Bst2, which acts as a restriction factor that prevents the release of enveloped viruses from infected cells by physically tethering the viral particles to the cell surface (S. X. Li et al., 2014; Mahauad-Fernandez et al., 2015). In addition, Bst2 has been recently described as a novel inhibitory receptor in NK cells, that is upregulated upon NK cell activation and is involved limiting their cytotoxic activity (Oh et al., 2022). In our analysis, Bst2 is most upregulated in Cluster 2, which is mainly composed of non-responding cMemory NK cells. It would be interesting to investigate whether Bst2 or Ly6A and Ly6E are directly involved in the recognition of *S. pneumoniae* by NK cells.

Overall, our results show that the effectiveness of cMemory NK cells does not rely in a small subset mounting an exceptionally strong response, but rather in a more generalised action that is supported by a moderate increase in the cytotoxicity across all cMemory NK cells combined with enhanced residence, survival and proliferative capacities (**Supplementary** Figure 7D). The fact that cMemory NK cells have a broad responsive phenotype yet less intense than the responding NK cells in cluster 1 is particularly interesting as several studies have described how, upon bacterial infections, NK cells can be deleterious, leading to tissue damage associated with an excessive inflammatory response (Christaki et al., 2015; Goldmann et al., 2005; Kerr et al., 2005; Wang et al., 2018). The strength of cMemory NK cells may lie in their capacity for efficient and widespread activation, rather than in an overly aggressive and potentially deleterious immune response. In conclusion, our findings define the transcriptional program of NK cells in the context of *S. pneumoniae* infection and therefore significantly advance our understanding the role of these cells during bacterial infections, thereby opening avenues for harnessing the potential of innate immune memory for therapeutic applications.

## MATERIALS AND METHODS

### Ethics statement

All protocols for animal experiments were reviewed and approved by the CETEA (Comité d’Ethique pour l’Expérimentation Animale - Ethics Committee for Animal Experimentation) of the Institut Pasteur under approval number dap17005 and dap230036, and were performed in accordance with national laws and institutional guidelines for animal care and use.

### Animal model

Experiments were conducted using C57BL/6J females of 7 to 12 weeks of age. CD45.2 mice were purchased from Janvier Labs (France). CD45.1 mice were maintained in house at the Institut Pasteur animal facility.

### Bacterial inocula

As in Camarasa, Torné et al 2023, all experiments that include infection with *S. pneumoniae* were conducted with the luminescent serotype 4 TIGR4 strain (Ci49) a commonly used pathogenic serotype containing pAUL-A Tn*4001 luxABCDE* Km’, (Saleh et al 2014), obtained from Thomas Kohler, Universität Greifswald. Experimental starters were prepared from frozen master stocks plated on 5% Columbia blood agar plates and grown overnight at 37°C with 5% CO_2_ prior to outgrowth in Todd-Hewitt (BD) broth supplemented with 50mM HEPES (TH +H) and kanamycin (50 μg/ml), as previously described in (Connor et al 2021). Inocula were prepared from frozen experimental starters grown to midlog phase in TH+H broth supplement with kanamycin (50 μg/ml) at 37°C with 5% CO_2_ in unclosed falcon tubes. Bacteria were pelleted at 1500xg for 10 min at room temperature, washed three times in DPBS and resuspended in DPBS at the desired CFU/ml. Bacterial CFU enumeration was determined by serial dilution plating on 5% Columbia blood agar plates.

For bacteria killed with paraformaldehyde (PFA), the concentrated bacteria, prior to dilution, were incubated in 4% PFA for 30 minutes at room temperature, washed three times in DPBS and diluted to the desired CFU/ml.

### Mouse infection

Animals were anaesthetized with a ketamine and xylazine cocktail (intra-peritoneal injection) prior to infection. Mice were infected with *S. pneumoniae* by intranasal instillation of 20μl containing 5x10^5^ (sublethal dose) or 1x10^7^ CFU (lethal dose). Control mice received intranasal injection of 20μl of DPBS. Animals were monitored daily for the first 5 days and then weekly for the duration of the experiment. Mouse sickness score from (Stark et al., 2018) was used to assess progression of mice throughout *S*. *pneumoniae* infection, representing the following scores: 0-Healthy; 1-transiently reduced response/ slightly ruffled coat/transient ocular discharge/up to 10% weight loss; 2 and 3-(2-up to 2 signs, 3-up to 3 signs) clear piloerection/intermittent hunched posture/persistent oculo-nasal dis-charge/persistently reduced response/intermittent abnormal breathing/up to 15% weight loss; 4-death. Animals were euthanized by CO_2_ asphyxiation at 20% loss of initial weight or at persistent clinical score 3.

### Sampling

Blood was collected by cardiac puncture at left ventricle with 20G needle and immediately mixed with 100mM EDTA to prevent coagulation. Spleens were disrupted by mechanical dissociation using curved needles to obtain splenocytes suspension in DPBS supplemented with 0,5% FCS+0,4% EDTA. Lungs were mechanically dissociated with gentleMACS Dissociator (Miltenyi Biotec) in cTubes containing lung dissociation kit reagents (DNAse and collagenase ref no. 130-095-927, Miltenyi Biotec).

### NK cell purification and transfer

NK cells were purified as described in Camarasa, Torné et al 2023. Splenocytes were passed successively through 100μm, 70μm and 30μm strainers (Miltenyi Biotec) in DPBS (+0,5% FCS+0,4% EDTA) and counted for downstream applications. NK cells were first enriched from splenocytes suspensions using negative enrichment kit (Invitrogen, ref no. 8804–6828) according to manufacturer’s protocol and combined with separation over magnetic columns (LS columns, Miltenyi Biotec). NK cells were then re-purified by negative selection using purification kit and magnetic columns according to manufacturer’s instructions to reach purities of approximately 98% (Miltenyi Biotec, MS columns and isolation kit ref no. 130-115-818). Purified NK cells were transferred intravenously by retro-orbital injections into recipient mice (0,5x10^6^ cells/mouse) in 100μl of DPBS or cultured for *in vitro* experiments.

### NK cell culture

Purified NK cells were cultured at 0.1x10^6^ cells/ml in 200μl RPMI 1640 (Gibco) supplemented with 10% FCS 1 U/mL Penicillin-Streptomycin (10 000 U/mL, Gibco) in 96 well round-bottom plates. Cells were either left unstimulated or put in contact with paraformaldehyde inactivated *S*. *pneumoniae* at a ratio of 40:1 and a cytokine cocktail composed of IL-15 (2 ng/ml), IL-18 (1,5 ng/ml) and IL-2 (1000 units/ml) (Miltenyi Biotec). After incubation at 37°C, cells were collected for protein staining and flow cytometry analysis. For proliferation assessment experiments, NK cells were stained with CFSE (Invitrogen) 2,5 mM in PBS for 5 minutes, before blocking in FCS and extensively washed with PBS before *in vitro* culture. For Killing assays, live *S. pneumoniae* was incubated in ½ diluted supernatants at an initial concentration of 500 CFU/ml, in a total volume of 120 ul. Bacterial CFU enumeration was determined by plating on 5% Columbia blood agar plates.

### Cell preparation for cytometry staining

Following mechanical dissociation, lung and spleen homogenates were filtered through 100μm strainers in DPBS supplemented with 0,5% FCS+0,4% EDTA, lysed in 1X red blood cell lysis buffer for 3 minutes (ref no. 420301, BioLegend), and passed successively through 70μm and 30μm strainers (Miltenyi Biotec). Single cell suspensions from lung and spleen were counted and prepared for surface staining in 96 well plates. For all, cells were first stained with anti-mouse CD16/CD32 to block unspecific binding and a cocktail of surface labeling antibodies for 40 minutes in DPBS (+0,5% FCS+0,4% EDTA). Next, cells were stained for viability using fixable viability dye (eFluor780, ref no. 65-0865-14, Invitrogen) for 5 minutes at 4°C and then fixed using commercial fixation buffer for 3 minutes (ref no. 420801, BioLegend). For intracellular staining of Granzyme B and Perforin, cells were permeabilized with a Fixation/Permeabilization commercial kit for 30 minutes (Concentrate and Diluent, ref no. 00-5123-43, Invitrogen). After permeabilization and wash, cells were stained with antibodies in DPBS (+0,5% FCS +0,4% EDTA) overnight at 4°C. After a final wash suspended in DPBS, sample acquisitions were performed on MACSQuant (Miltenyi Biotec) flow-cytometer and analyses were done using FlowJo Software (TreeStar).

### Antibodies

CD45 (30-F11): Termo Fisher Scientific, Catalog #11-0451-82. CD45.1 (A20): BioLegend, Catalog #110741. CD45.2 (104): Termo Fisher Scientific, Catalog #12-0454-82. NK1.1 (PK136): BioLegend, Catalog #108732. CD3 (145-2C11): Termo Fisher Scientific Catalog #25-0031-82. Ly6ae (D7): Termo Fisher Scientific, Catalog #12-5981-82. CD69 (H1.2F3): Termo Fisher Scientific, Catalog #12-0691-83. CD49a: BD Biosciences, Catalog #562115 (Ha31/8 (RUO)). CD49b (DX5): Termo. Fisher Scientific, Catalog #17-5971-82. BIM (C34C5): Cell Signalling technologies, Catalog #12186S. BCL2 (10C4): Termo Fisher Scientific, Catalog #11-6992-42. GzmB (GB11): BioLegend, Catalog #515406. Perforin (S16009A): BioLegend, Catalog #154302. Ki67 (16A8), Sony. Catalog #3862030.

#### 10x Library preparation, sequencing and alignment

ScRNA-seq libraries were generated with the Chromium Single Cell 3ʹ v.2 (Day 7) and v.3 (Day 14) assay (10x Genomics). Libraries were sequenced using the HighSeq 4000 platform (Illumina) to a depth of about 300 million reads per library with 2 × 50 read length. Raw reads were aligned to mouse genome (mm10) and cells were called using cellranger count (v.3.0.2).

#### scRNA-seq analysis

Single cell was analysed using the R package Seurat v5.1.0 (Hao et al., 2024). A set of 4918 high quality cells (2453 from cc sample and 2465 cMemory sample) were chosen as those with 1000 to 3500 genes detected and less than 20% mitochondrial gene expression. Additionally, ribosomal genes were filter out, leaving 32184 genes for the downstream analysis. The dataset was then normalized, mitochondrial gene was regressed and variable features selected expression using the sctransform method as implemented in Seurat (Choudhary & Satija, 2022). Linear dimension reduction was performed by principal component analysis (PCA) and these components were used to merge cNaïve and cMemory samples by reciprocal PCA method (RPCA). This last embedding was used to build the shared nearest-neighbor graph with 40 RPCA component. Cells were then clustered using Seurat’s implementation of the Louvain algorithm and a resolution of 0.3. For the purpose of visualization, the nonlinear dimensionality reduction UMAP was calculated with default parameters over the RPCA components. Visualization of the expression of specific genes or groups of genes was done with the Single Cell sHiNy APPlication (SCHNAPPs) (Jagla et al., 2021).

#### Cluster Functional analysis

To characterize functionally each cluster, gene markers conserved among cNaïve and cMemory conditions were calculated by performing differential gene expression testing for each condition and combining the p-values using meta-analysis methods from the MetaDE R package. We then performed a Gene Set Enrichment Analysis (GSEA) (Subramanian et al., 2005) for the 4966 marker genes common to all clusters, by ranking them using the average log fold change from the differential gene expression above. We used the gene sets from the Reactome database and the GSEA implementation in https://gitlab.pasteur.fr/hvaret/fgsa_scripts.

#### Differential analysis

Differences between cMemory and cNaïve conditions were estimated by adapting a differential state analysis at a pseudo-bulk as described in (Crowell et al., 2020). Briefly, cells in each cluster were subclustered as described above, but using 50 RPCA components for the nearest-neighbor graph and resolution of 1 for the clustering. These subclusters represent the transcriptional variability with each cluster and are treated as replicates. A standard genes x samples count matrix is defined per cluster, by summing the cMemory or cNaïve per subcluster for each gene. Lowly expressed genes are filtered and only the 2907 genes with more than 200 counts over all samples are kept. Differential analysis between cMemory and cNaïve is performed per cluster, following the standard procedure for RNA-seq with limma, adding the replicate as a covariate of the linear model. Gene sets enriched among cMemory or cNaïve upregulated genes were calculated by applying GSEA approach described above and using the differential analysis results of each cluster over the 2907 filtered genes.

#### Cell trajectories analysis

Cell trajectories were inferred on the cMemory and cNaïve datasets, projected over the RPCA UMAP embedding with the R package slingshot (Street et al., 2018). A pseudotime variable is estimated for cells along each trajectory, which reflects its relative position. We set the connection distance (i.e. omega) to the median edge length of the unsupervised minimum spanning tree scaled by the default value 3, thus allowing for multiple trajectories. The central cluster 0 was set as the starting point for the construction of the minimum spanning tree(s). To identify the genes whose expression is significantly associated with the inferred cMemory and cNaïve lineages, i.e. the driving genes, the expression of the 3e3 more variable was regressed over the inferred cell trajectories using a general additive model (GAM) and a negative binomial noise distribution as implemented in the tradeSeq R package (Van den Berge et al., 2020). Significance of the association was then calculated using the association test of the same package.

## Data and code availability

The scRNA-seq raw data reported in this paper are available in GEO database under the accession number GSE208362. https://www.ncbi.nlm.nih.gov/geo/query/acc.cgi?acc=GSE208362

## ACKNOWLEDGEMENTS

We would like to thank the members of Chromatin and Infection Laboratory for valuable input throughout the project and for critical reading of the manuscript. We would also like to thank the members of the Hub of Bioinformatics and Biostatistics for useful discussions. This work used the computational and storage services (MAESTRO cluster) provided by the IT department at Institut Pasteur, Paris. We acknowledge the staff of the breeding zone in the CFJ animal facility (BIME_EOPS) of C2RA, Institut Pasteur, to produce mice for the *in vivo* experiment. This project was supported by the Institut Pasteur, the Agence Nationale de la Recherche (ANR-17 CE120007 01 EPIBACTIN), the Fondation pour la Recherche Médicale (FRM608 EQU202003010152), the Fondation iXCore-iXLife, the Don Prix CANETTI 2020, the EMBO Young Investigator Program. M.A.H. is a member of the Laboratoire d’Excellence “Integrative Biology of Emerging Infectious Diseases” Agence Nationale de la Recherche (ANR-10-LABX-62-EIBID). J.T was supported by the Fondation ARC Cancer Research (grant ARCPOST-DOC2021070004074). T.M.N.C received a salary from the Crédit Agricole d’Ile de France and Fondation pour la Recherche Médicale, grants no. PMJ201810007628 (Prix MARIANE JOSSO) and no. FDT202106012790 (Fin de thèse). The Bioinformatics and Biostatistics Hub is supported by Institut Pasteur.

---

**Supplementary Figure 1.** NK cells purification for transfer and isolation. **A)** Representative gating strategy to measure purity of transferred D21PBS NKs or D21SPN NKs before transfer to recipient mice. Purity represents the percentage of NK cells among CD45.2+ transferred cells. **B)** Following NK cells transfer (CD45.2+) and lethal infection of the recipient mice (CD45.1+), representative gating strategy to identify transferred NK cells (CD45.2+) or endogenous NK cells (CD45.1+) in the lungs of recipient mice at 40 hours post lethal infection. **C)** CFU counts in the lungs of mice having received D21PBS NK cells (black dots) or D21SPN NK cells (red dots) 40h post lethal infection. Box plot where each dot represents an individual mouse, lines are the mean, error bars show min to max. **D)** Cell distribution among clusters obtained using different values of resolution (i.e. 0.3, 0.6 and 1.0) for the clustering algorithm. Bar plots below represent the corresponding proportion of cNaïve and cMemory cells per cluster. (**E)** Number of NK cells per cluster.

**Supplementary Figure 2.** Characterizing the functional diversity of NK cell clusters 3,4 and 5. **A)** Dot plot representing the expression of top 9 marker genes, conserved among cNaïve and cMemory cells and ranked by their average log fold change, for clusters 3, 4 and 5. The size of the dots represents the percentage of cells expressing the corresponding gene and the colour represents its average expression. **B)** Classification of marker genes in encoding membrane markers, secreted proteins, or transcription factors (TF). **C)** Gene Set Enrichment Analysis (GSEA) for each set of markers conserved among cNaïve and cMemory in every cluster. Selected REACTOME gene sets significantly enriched (adjusted p-value < 0.05) among upregulated or downregulated genes in clusters 3, 4 and 5 are compared using the Normalized Enrichment Score (NES). N=Number of genes in every gene set.

**Supplementary Figure 3.** Ifngr1 is downregulated in cMemory NK cells. Violin plots representing the sum of the normalize gene counts for *Ifng* **(top panel)** or Ifngr1 **(low panel)** in the cMemory or cNaïve population within every cluster.

**Supplementary Figure 4.** Top driving genes of single cell trajectories. A-B) Upset plots comparing the top driving genes for trajectories calculated over cNaïve and cMemory NK cells, starting in Cluster 0 **(A)** or Cluster 2 **(B)**. Top 100 significant driving genes of each trajectory were used for this analysis and only comparisons including 5 or more genes are shown. **C)** Over-representation of biological processes among the top driving genes of the above-mentioned trajectories.

**Supplementary Figure 5.** cMemory NK cells have a more active phenotype across clusters. A-B) Violin plots representing the sum of the normalize counts per gene in the cMemory or cNaïve population within every cluster for **(A)** *Cd69*, **(B)** *Ly6a or Ly6e*. **C)** Combinations of normalized counts for *Cd69, Ly6a* and *Ly6e* in the total cNaïve or cMemory populations. **D, E)** C57BL/6 mice were intranasally infected with either PBS (black symbols) or sub-lethal dose of S. After 21 days, NK cells were highly purified from spleens of D21PBS or D21SPN mice and transferred to recipient mice. **D)** Flow cytometry analysis of the percentage of CD69+ NK cells **(top panel)** or Ly6ae+ NK cells **(low panel)** among transferred (CD45.2+) cNaïve or cMemory NK cells in spleen and its MFI for Cd69 or Ly6a. **E)** Flow cytometry analysis of the percentage of CD69+ NK cells **(top panel)** or Ly6ae+ NK cells **(low panel)** among endogenous (CD45.1+) NK cells in lung or spleen from mice having received cNaïve or cMemory NK cells and its MFI. **F)** Flow cytometry analysis of the absolute number of NK cells in lung and spleen of mice having received Memory or Naïve NK cells. **G)** Flow cytometry analysis of the percentage of Ki67+ NK cells in lung or spleen from mice having received cNaïve or cMemory NK cells and its MFI. Each dot in the boxplots represents an individual mouse (black dots for mice having received D21PBS NKs and red dots for mice having received D21SPN NKs), lines are the median and error bar show min to max. Data are pooled from n≥3 repeats with n≥3 mice/group. ns, not significant. * p < 0.05 and ** p <0.01, t-test with crossed design.

**Supplementary Figure 6.** : cMemory NK cells exhibit enhanced residency and survival phenotype. **A,B,E)** C57BL/6 mice were intranasally infected with either PBS (black symbols) or sub-lethal dose of *S. pneumoniae* After 21 days, NK cells were highly purified from spleens of D21PBS or D21SPN mice and transferred to recipient mice. **A)** Flow cytometry analysis of the percentage of Cd49b+ NK cells **(top panel)** or Cd49a+ NK cells **(low panel)** among endogenous (CD45.1+) cNaïve or cMemory NK cells in blood (**left**) or lung **(right**) and its MFI for DX5 or CD49a. **B)** Flow cytometry analysis of the percentage of Cd49+ NK cells **(top panel)** or Cd49a+ NK cells **(low panel)** among transferred (CD45.2+) or endogenous (CD45.1+) cNaïve or cMemory NK cells in Spleen its MFI for DX5 or CD49a. **C)** Violin plots representing the sum of the normalised read counts for *Bcl2l11* (Bim) **(top panel)** or *Bcl2* **(low panel)**, in the total of cNaïve or cMemory NK cells. **D)** Violin plots representing the sum of the normalize gene counts for *Bcl2l11* (Bim) **(top panel)** or Bcl2 **(low panel)** in the cMemory or cNaïve population within every cluster. **E)** Flow cytometry analysis of the percentage of Bim+ NK cells or Bcl2+ NK cells among endogenous (CD45.1+) cNaïve or cMemory NK cells in spleen **(top)** or lung **(bottom)** and MFI for Bim or Bcl2. Each dot in the boxplots represents an individual mouse (black dots for mice having received D21PBS NKs and red dots for mice having received D21SPN NKs), lines are the median and error bar show min to max. Data are pooled from ≥ 3 repeats with n≥3 mice/group. ns, not significant. * p < 0.05 and ** p <0.01, p < 0.001, t-test with crossed design.

**Supplementary Figure 7.** : cMemory NK cells have a more responsive phenotype across clusters. A-B) C57BL/6 mice were intranasally infected with either PBS (black symbols) or sub-lethal dose of S. pneumoniae After 21 days, NK cells were highly purified from spleens of D21PBS or D21SPN mice and transferred to recipient mice. A) Flow cytometry analysis of the percentage of GzmB+ NK cells (top) or Perf+ NK cells (bottom) among transferred (CD45.2+) cNaïve or cMemory NK cells in spleen and MFI. B) Flow cytometry analysis of the percentage of GzmB+ NK cells (top) or Perf+ NK cells (bottom) among endogenous (CD45.1+) cNaïve or cMemory NK cells in lung (left) or spleen (right) and MFI. C) Bacterial counts for live S. pneumoniae incubated for 4h in ½ diluted supernatants from NK cells previously incubated with cytokines and with or without FISP. D) Memory was induced in donor mice (CD45.2), and their cells were transferred into CD45.1 recipient mice, which were then challenged. scRNA-seq of sorted CD45.2 cells revealed distinct NK cell subpopulations, including a highly responsive cluster containing both cMemory and cNaive NK cells and a cluster of non-responsive cMemory NK cells. In the remaining clusters, cMemory NK cells exhibited enhanced response features compared with cNaive NK cells. FACS analysis, guided by the scRNA-seq data, confirmed that cMemory NK cells displayed increased cytotoxicity, activation, survival, residency, and proliferation compared with cNaive NK cells. Each dot in the boxplots represents an individual mouse (black dots for mice having received D21PBS NKs and red dots for mice having received D21SPN NKs), lines are the median and error bar show min to max. Data are pooled from ≥ 3 repeats with n≥3 mice/group. ns, not significant. * p < 0.05 and ** p <0.01, p < 0.001, t-test with crossed design.

**Supplementary Table 1:** Differential state analysis results. Gene statistics are shown for the comparison of cMemory vs cNaïve cells for every cluster.

**Supplementary Table 2:** Gene set enrichment analysis results. Gene set statistics are shown for gene sets upregulated in cMemory vs cNaive cells for every cluster.

