## supplementary material for "NK cells undergo transcriptional and functional reprogramming following *Streptococcus pneumoniae* infection"

**Supplementary Figure 1**. NK cells purification for transfer and isolation.

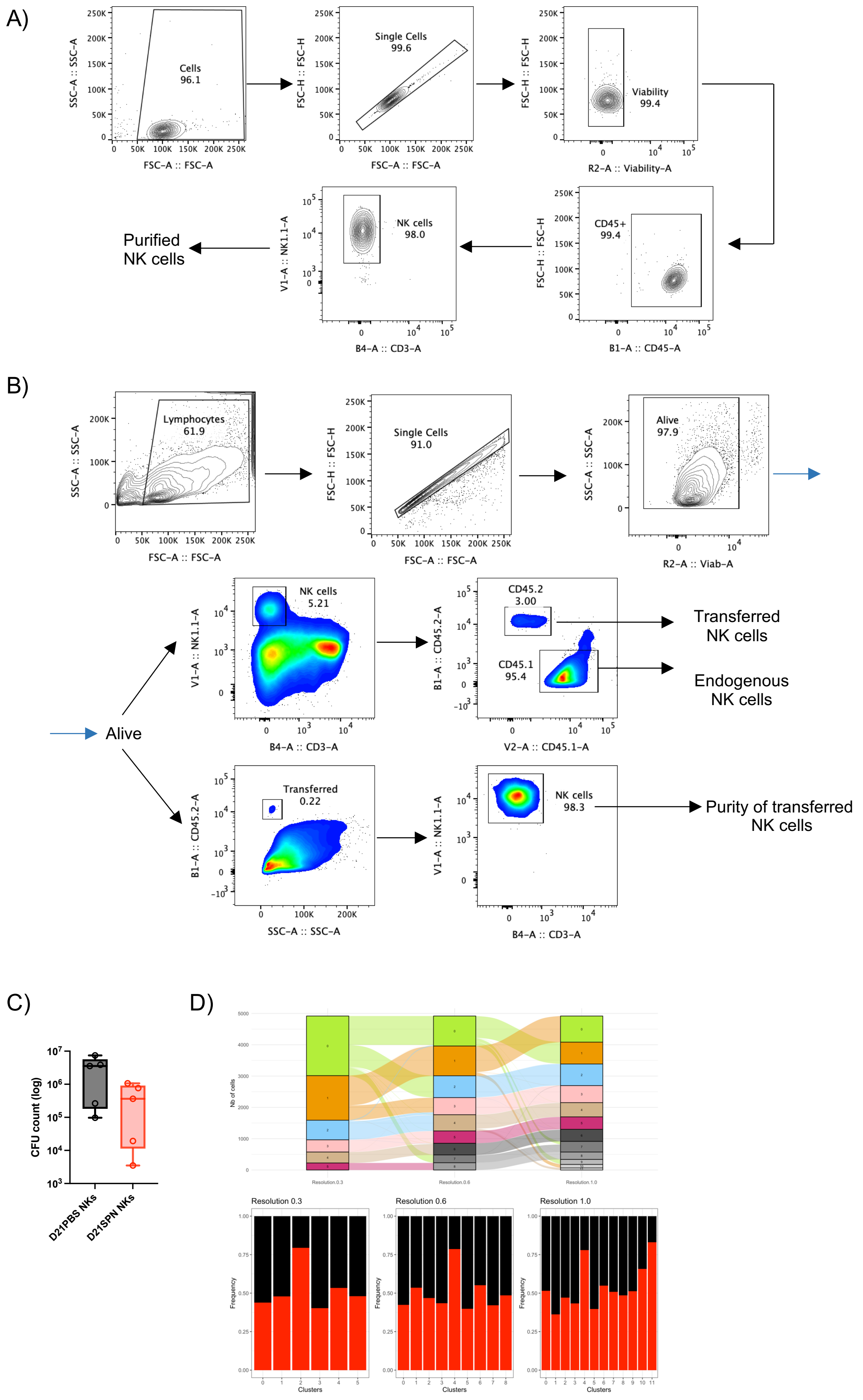

**Supplementary Figure 2**. Characterizing the functional diversity of NK cell clusters 3,4 and 5.

A)

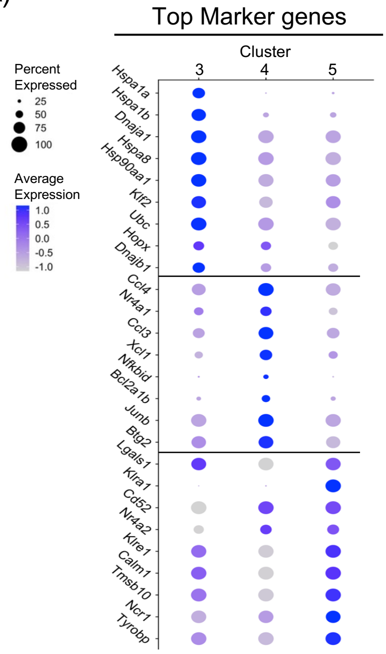


B)

| | MEMBRANE | SECRETED | TF |
|---|---|---|---|
| **3** | Klra9<br>Klrg1 | Ctsd<br>Ctsc<br>Cma1 | Klf2 |
| **4** | Cd9<br>Icam1 | Ccl3<br>Ccl4 | Junb<br>Nfkbid<br>Nr4a1<br>Egr3 |
| **5** | Klra1 (Ly49a)<br>Klre1<br>Calm1 | Ctsw | Nr4a2 |

C)

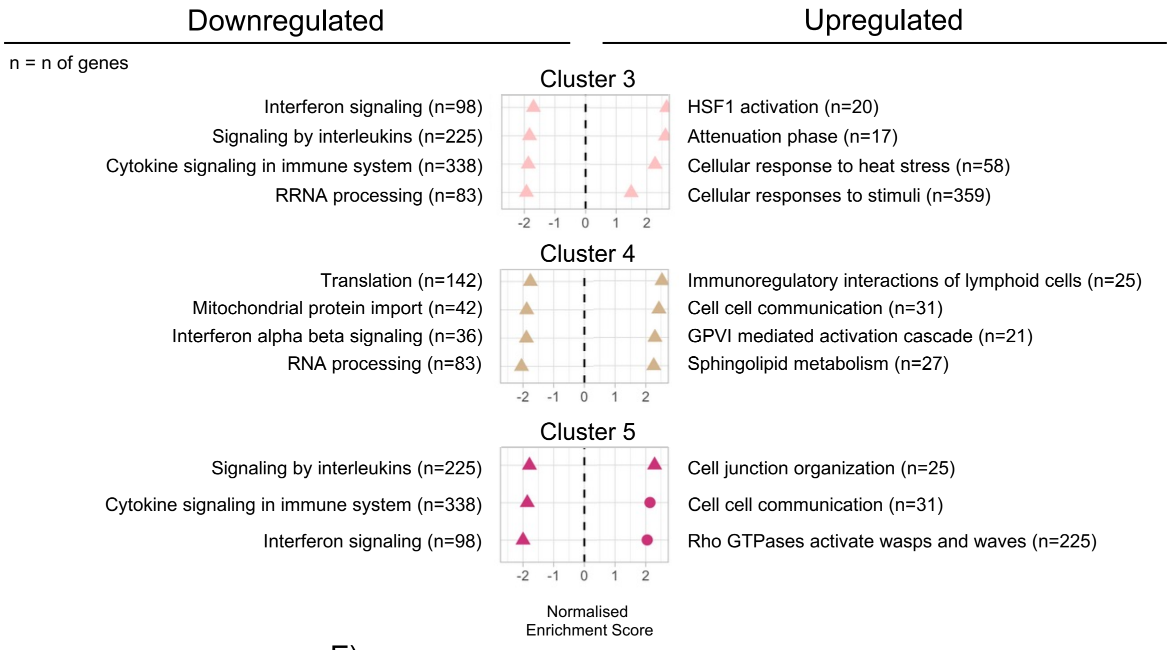


D)

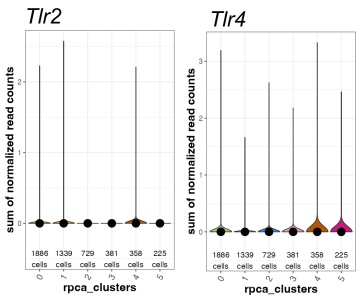


E)

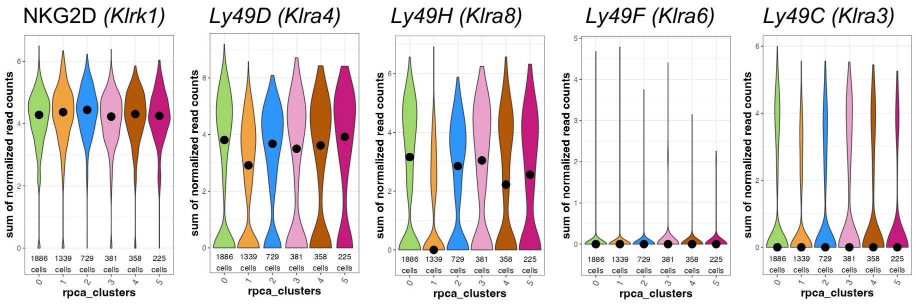

**Supplementary Figure 3**. Ifngr1 is downregulated in cMemory NK cells.

A)

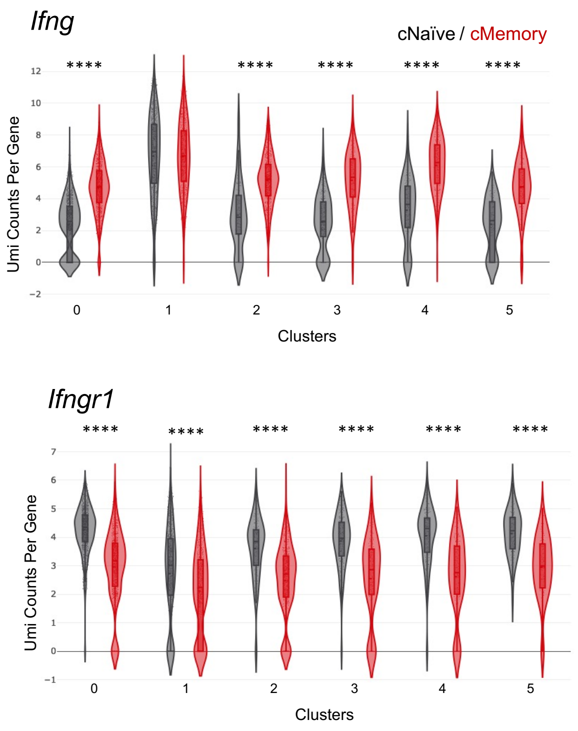

**Supplementary Figure 4**. Top driving genes of single cell trajectories.

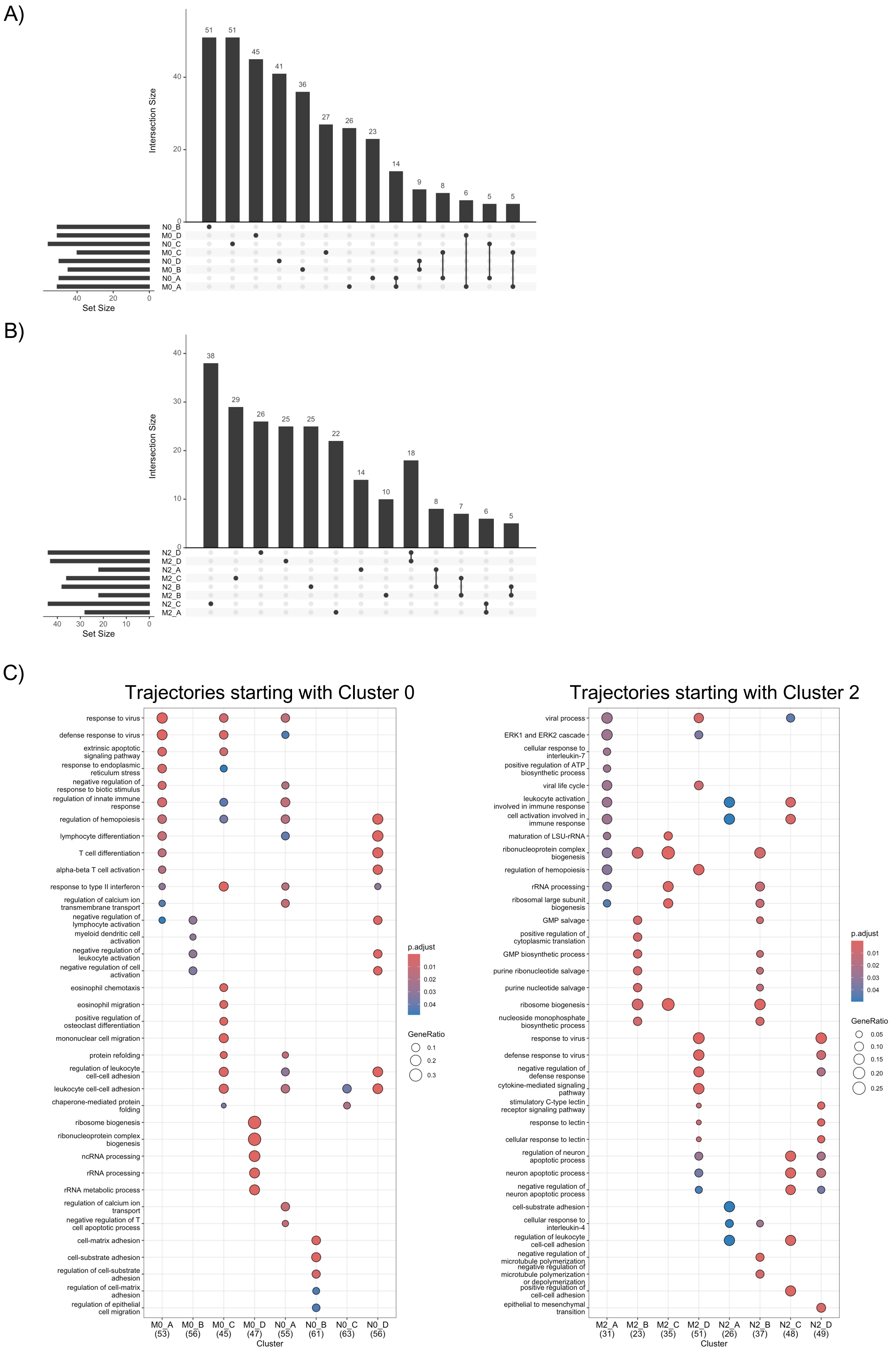

**Supplementary Figure 5**. cMemory NK cells have a more active phenotype across clusters.

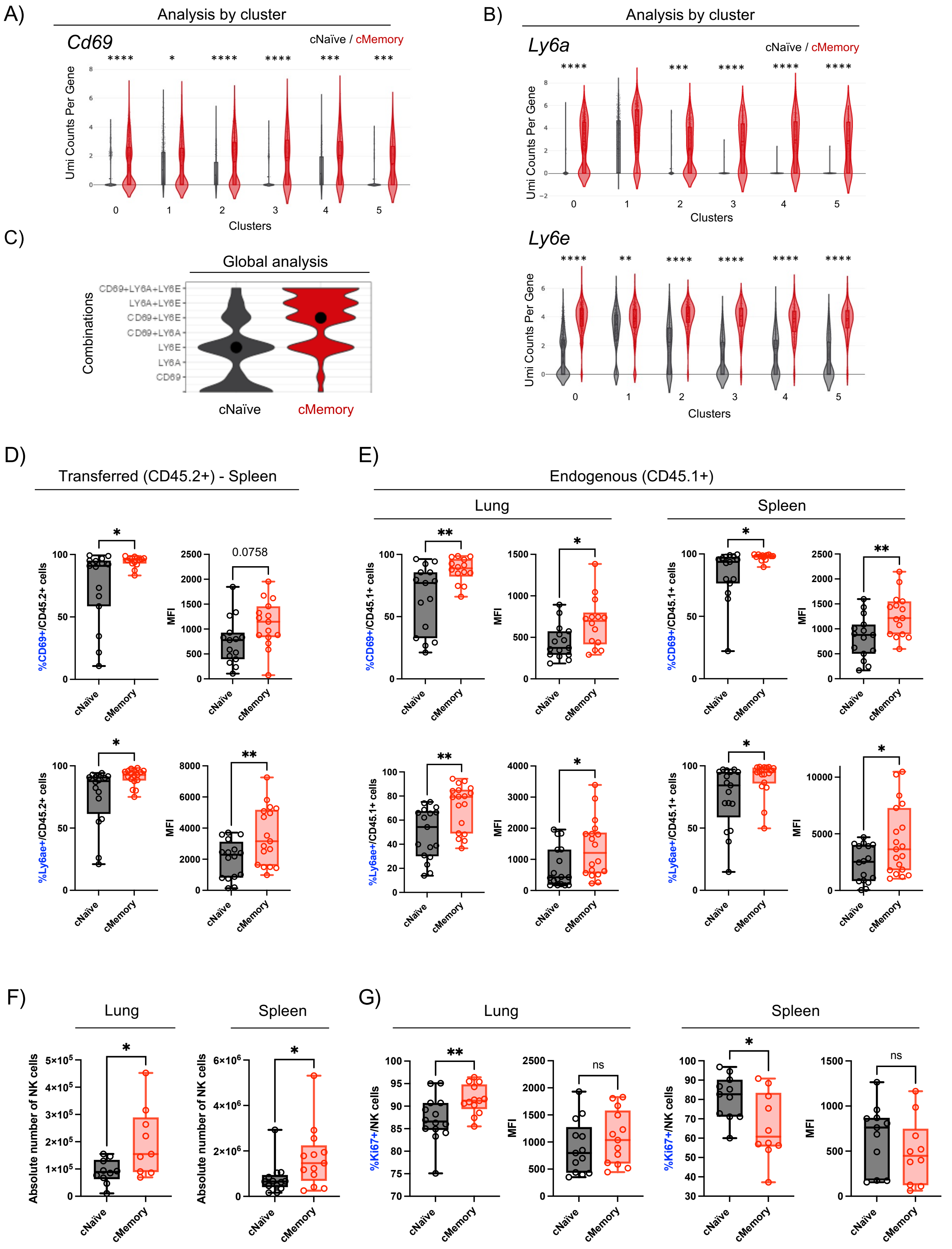

**Supplementary Figure 6.** cMemory NK cells exhibit enhanced residency and survival phenotype.

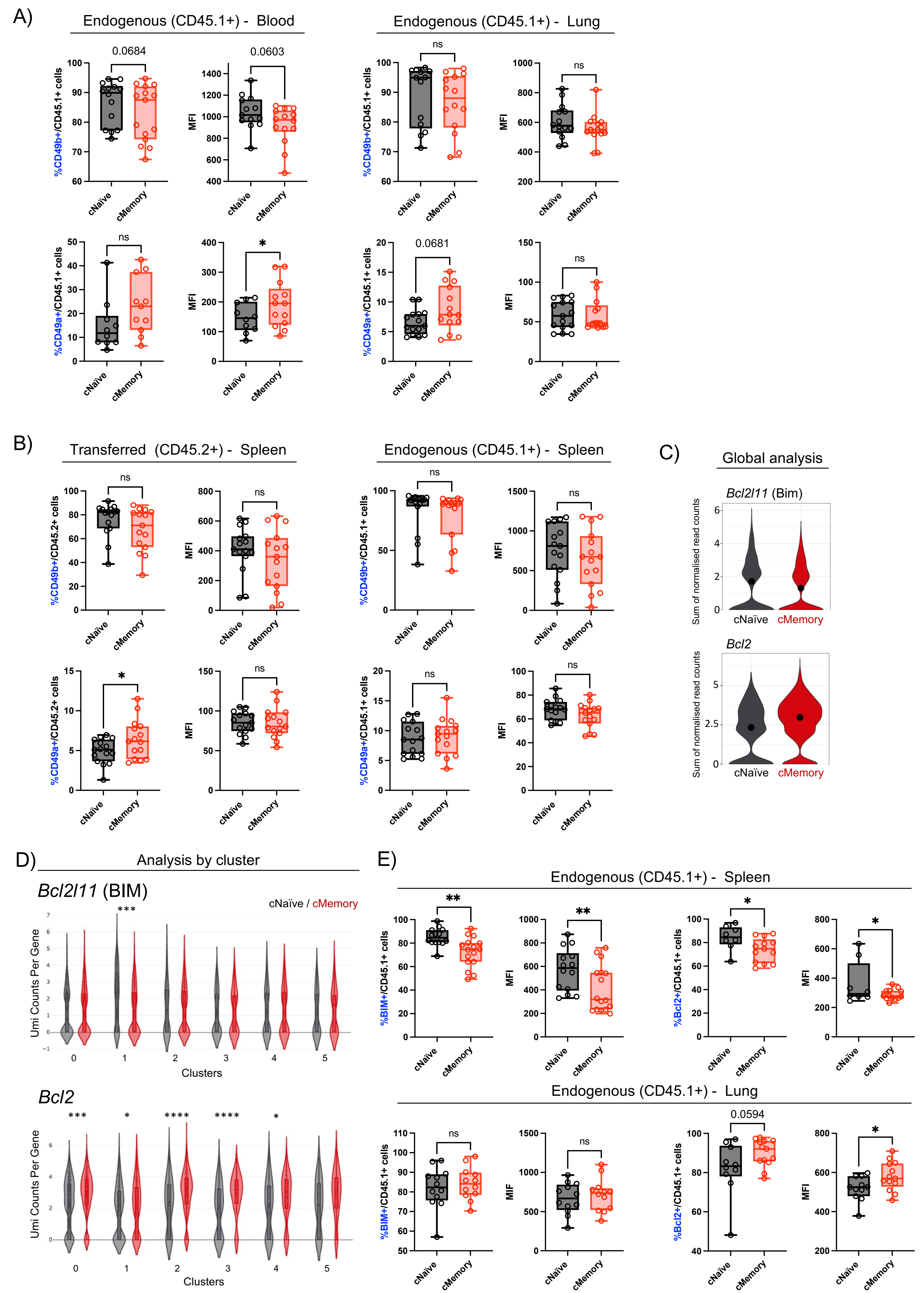

**Supplementary Figure 7.** Memory NK cells have a more responsive phenotype across clusters.

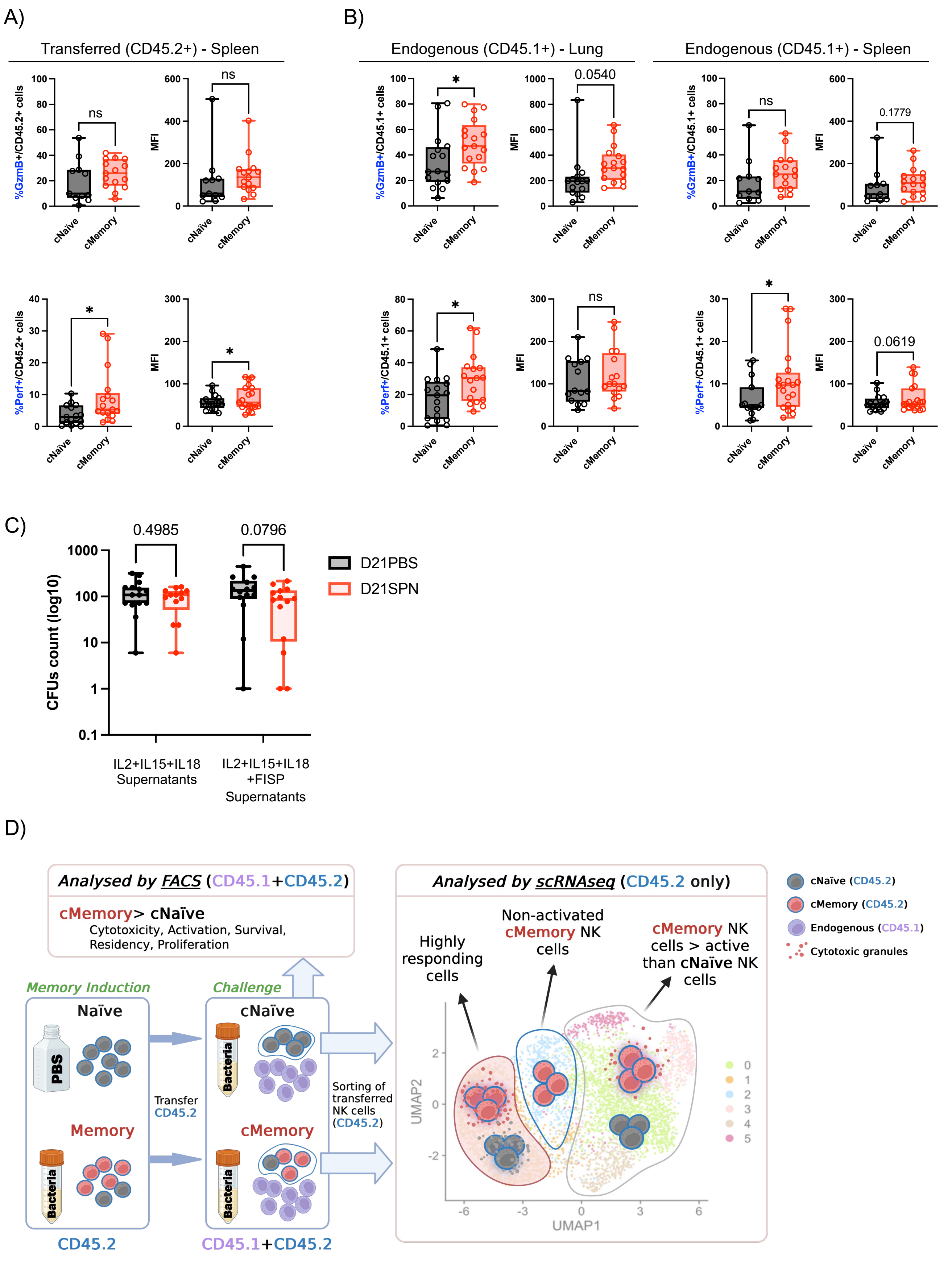

**Table 1:** Differential state analysis: Genes upregulated in a specific cluster

| Clusters | Downregulated in cMEMORY |
|---|---|
| 0 | Apbb1ip, Egln1, Kdm7a, Mat2b, Mkrn1, Sike1, Slc20a1, Ugcg |
| 1 | 1810041H14Rik, Egln2, Gimap5, Gm49980, Gna13, H2-Q4, Irak2, Plscr1, Prr7, Tet2, Zeb1 |
| 2 | Bri3, Cx3cr1, Cxcr4, Fosb, Ifnar1, Il4ra, Klrg1, Lncpint, Nfkbid, Ptger4, R3hdm4, Rel, Zfpm1 |
| 3 | 0610012G03Rik, As3mt, Atp6v0a2, Ccnt1, Cers4, Ctsd, Cwc25, Elmo1, Gfod1, Hist1h1c, Itga4, Itgb2, Lysmd3, Maz, Necap2, Ppm1a, Prkacb, Prpf19, Ptpn7, Pycard, Rbbp8, Sephs2, Sfr1, Smc6, Spata13 |
| 4 | 4930523C07Rik, Avl9, Bbc3, Bex3, Ccni, Efr3a, Fam53b, Fmnl1, Gatad2b, Ggct, Haus3, Hook3, Kdm2b, Klf3, Kmt2d, Mcm6, Me2, Mkln1, Pcm1, Pde2a, Pde7a, Phlda1, Pik3cd, Pnrc1, Ppp1r16b, Ralbp1, Rcc2, Rnf138, Selenoh, Sema4a, Sept6, Smarcc1, Sptssa, St8sia4, Stk10, Stk11, Sun2, Tlk1, Tmem123, Tnfaip3, Tnrc6b, Tpi1, Trbc2, Usp34, Zfp326 |
| 5 | Brd1, Cd27, Cyth1, Evl, Fam32a, Hnrnpul1, Irf2bpl, Klf7, Kmt2c, Lasp1, Nelfb, Orai1, Pitpnc1, Ppig, Saraf, Sdhd, Sft2d1, Smad7, Spn, Ssna1, Tagapm Tmem234, Tnks2, Ube2r2 |
| **Clusters** | **Upregulated in cMEMORY** |
| 0 | Casp8, Cdkn1a, Hopx, Hspa5, Il2ra, Mettl1, Mir155hg, Mxd1, Myd88, Pim1, Pim2, Plscr1, Stk39, Stom |
| 1 | Cpne3, Hsp90b1, Hsph1, Klra1, Klrb1f, Klrd1, Klre1, Pcbd2, Ptpn6, Rnaset2b, Styk1 |
| 2 | Daxx, Samd9l |
| 3 | Agfg1, Atf6, Bcap29, Capg, Ccl4, Cnp, Ctla2a, Emb, Fndc3a, Gimap1os, Gm2000, Gpr171, H13, Lars, Lpp, Ms4a4b, Myo6, Naa20, Prrc1, Psmc1, Snrnp27, Zswim6 |
| 4 | Batf, Gfpt1, Lgals9, Mthfd2, Prf1, Sec61b, Selenos, Spcs3, Xbp1 |
| 5 | Arl5a, Atp2a2, Cx3cr1, Irf1, Mylip, Nmt1, Pdia4, Psma5, Relb, Ruvbl1, Ssr1 |

**Table 2**: Differential state analysis: Genes upregulated in an intersection of clusters

| Clusters | Downregulated in cMEMORY |
|---|---|
| **2-3** | Itgam, Klrb1b, Lgals1, Ptms, S100a6, Sgk1, Sipa1, Timp2, Zmiz1 |
| **4-5** | Coq10b, Galnt2, Gsap, Kpna4, Limd1, Ripor2, Rp9, Slc25a36 |
| **3-4** | Ankrd13a, Cdkn2aip, Hsph1, Prex1, Sppl3, Topors, Trdc |
| **3-5** | Bin1, Dnajb4, Fli1, Myl12b, Psap |
| **2-3-4** | AB124611, Dbf4, Dgkz, Gm45716, Lsp1, Tnik, Ubash3b |
| **0-2-4** | Bambi, Fosl2, Lamp1, Myb, Sik, Tob1 |
| **0-3-4** | Maf1, Pbxip1, Pgp, Pik3r1, Rassf1, Yaf2 |
| **0-3-4-5** | Bin2, Cbx4, Cd2, Ifngr1, Ing2, Klhl24, Rnf166, S100a10, Supt4a, Tprgl, Usp48, Vps37b, Zbtb20 |
| **0-2-4-5** | Hmgb2, Hspa1a, Hspa1b, Rb1cc1, Rpa2, Tnfsf12 |
| **0-2-3-4** | Arhgap31, Hhex , Rap2b, Rnf130, S1pr4 |
| **0-2-3-4-5** | Add3, Adgre5, B4galt1, Bnip3l, Cd7, Cd72, Cd9, Cks2, Cox7a2l, Crip1, Fam89b, Gpc1, H2afv, Itgb7, Jakmip1, Mxd4, Nr3c1, Nr4a1, Prkca, Rasgrp2, Rgs2, S1pr5, Samd3, Tbc1d10c, Tm6sf1, Tspan32, Vegfa, Ypel3, Zfp36l1 |
| **0-1-2-3-4-5** | Ccl5, Cma1, Gm10076, Gstp1, Spry2, Uba52 |
| **Clusters** | **Upregulated in cMEMORY** |
| **0-3** | 9930111J21Rik2, Apobec3, Dtx3l, Gbp9, Gm43305, Mapk6, Nmi, Picalm, Smchd1, Stx11, Tnfrsf9 |
| **0-4** | AU020206, Car5b, Cgas, Gbp5, Manf, Plac8, Rrp1b, Slc16a6, Slc35b1 |
| **0-3-4** | Aim2, B930036N10Rik, Bcl3, Cd53, Clec2d, Cysltr2, Flot1, Gbp2, Gbp4, Ifi211, Oasl1, Parp14, Rpn1, Sec61, Sell, Serpina3f, Vars, Zbtb32 |
| **0-3-5** | Dgat1, Gadd45b, H2-T23, Mndal, Nfkbiz, Psme1, Rhoh, Ssrp1, Trafd1 |
| **0-3-4-5** | 5830428M24Rik, Calr, Casp4, Ccr5, Clic4, Creld2, Csrnp1, Cxcl10, Frmd4b, Furin, Gpr155, Gpr18, Gzmb, Herc6, Ifi204, Ifi35, Irf7, Isg20, Lilrb4a, Mrpl52, Ms4a6b, P2ry14, Pdia3, Pdia6, Pml, Psme2, Rsad2, Sbno2, Serpina3g, Slamf7, Slfn8, Socs3, Tap1, Tspan13 |
| **0-2-3-4** | AA467197, Adrb2, Cish, Ier3, Il12rb1, Mctp2, Samhd1, Sult2b1 |
| **0-2-3-4-5** | Atf3, Bst2, Cd69, Ctss, Dgkh, Eif2ak2, Gadd45g, Gbp6, Gimap7, Gng2, Ifi203, Ifi206, Ifi208, Ifi209, Ifi213, Ifi214, Ifi47, Ifng, Isg15, Jaml, Lgals3bp, Lilr4b, Ly6a, Ly6e, Maff, Map3k8, Muc1, Parp9, Phf11a, Phf11b, Ppa1, Prdm1, Rnf213, Rras2, Rtp4, Socs1, Trim12c, Trim30a, Trim30d, Usp18, Xaf1, Xcl1, Zbp1 |

**Table 3**: Top driving genes of single cell trajectories starting in Cluster 0

| Trajectory | cNAÏVE |
|---|---|
| **N0_A** | Ly6a Furin Hspa1b Klra9 Sdc4 Klra4 Klra8 S100a4 Gem Ccl5 Vim Ncl Mir155hg Gm43305 Xist Satb1 Hif1a Mndal Gimap5 Nr4a2 Tspan13 Zfp36l2 Gbp5 Txnip 1810041H14Rik Neurl3 Spry2 Serpinb9 P2ry14 Dnaja1 Klrd1 Klrg1 Snx10 Slfn2 Sfi1 Tmsb10 mt-Atp6 Ahnak Cd7 Sec61g Herc6 Arid5b Ctsd Sytl3 Itgb7 Ptpn6 Ptma Cish Uqcrq Adgre5 Hint1 Parp14 Ier2 Atf6 Rilpl2 Cd53 Zeb2 Eif4ebp1 Hsp90aa1 Ptms |
| **N0_B** | Setdb2 Eef1a1 Plk3 Rnf43 Tbrg1 Sat1 Mdn1 Klra3 Gm26740 Sipa1l1 Atp5g2 Fubp1 Zyg11b Eef2 Usp10 Calr 4930581F22Rik Pim2 Mrpl12 Pgls Aprt Eml4 Atg16l2 Rab8b Pglyrp1 Hspd1 Tubb5 Itga6 S100a11 B9d2 Rbm3 Pitpnc1 Tef Ddx11 Gm10076 Irs2 Kdm3b 1110059G10Rik Wars Nf1 Gpx4 Itgax Keap1 Ttf1 Dapk2 Ninj1 Gar1 Nsun2 Cmtm7 Macf1 Gm17056 Lats2 AI504432 Map4k4 Setbp1 Ranbp1 Lsp1 Mthfd2 Lars Klra7 Tagln2 Tapbp Znfx1 Rnf213 Hsp90b1 Trpm7 Rnf216 |
| **N0_C** | Hspa1a S100a6 Mbtps2 Dgat1 Hsph1 Lgals1 Gzma Shisa5 Rpap3 Ifngr1 Ubc Cenpa Tmsb4x Ppp1r14b Pbrm1 Sp4 Sell Chd2 Cyba Pja1 Sub1 Stat4 Pmaip1 Uspl1 Larp7 Akr1b10 Tec Tm6sf1 Gna13 Ubqln2 Kctd12 Ddx27 Itgb2 Slc25a24 mt-Co2 Ssh2 Trp53i11 1110038B12Rik Myo1g Prf1 Sars Eef1b2 Bin2 Gas7 Ftl1 Kdm7a Gm36723 Chordc1 Tuba1b Bcl3 Fkbp2 H2-T22 Gmip Lax1 Pfkp Flna Tuba1a Smc6 Sla Gramd3 Dop1b Sptbn1 Lilr4b Tmem127 Adap1 Ly75 Dynll1 Mgat5 1700023H06Rik Slc25a5 |
| **N0_D** | Ccl4 Pim1 Nfkbid Jund Bcl2a1d Ier5 Nr4a3 Id2 Lfng Relb Zc3h12a Prr7 Tgif1 Fbl Kdm2b Rgs16 Slc38a1 Gpr65 Ppp1r16b Cd69 Nfkbiz Junb H2-Q6 Actg1 Btg2 Gtpbp4 Eef1e1 Apbb1ip Crtam Hivep2 Adora2a Btg1 Pde2a Limd2 Ubald2 Plek Nfkb2 Cd2 Zfp36 Ccnd3 Klf13 Klrb1f Ypel3 Phb Phlda1 Mxd4 Map3k8 Mapk6 Ly6e Dennd4a Hspa9 Egr3 Gapdh Actb Rasgrp2 Ran Atic |
| **Trajectory** | **cMEMORY** |
| **M0_A** | Ly6a Gzmb Isg15 Atf3 Il2ra Tgfb1 Traf1 Bst2 Nr4a2 Tspan13 Zfp36l2 1810041H14Rik Neurl3 Spry2 Klrd1 Bcl2 Zfp52 Vps37b Shisa5 Frmd4b Tmsb10 Junb 1110038B12Rik Sec61g Ccnd2 Tmsb15b2 Ncr1 Adgre5 Ier2 Atf6 Cd53 Snhg1 Ifngr1 Mettl1 Hopx Dnajb1 C1qbp Ifi47 H2-Q7 Ranbp1 Sipa1l1 Ptp4a2 Samd9l Gpatch4 Serinc3 Bcl2l1 Taf1d Gar1 Pik3r1 Adap1 Polr3d Tespa1 Pnrc1 |
| **M0_B** | Ggps1 Srgn Tubgcp2 Ddx21 Setbp1 Hccs Arhgap45 Gapdh Nr4a1 Txnip Rilpl2 Nolc1 Samsn1 Atp1b1 Gbp5 Pim3 Spag9 Bcl2a1d Gnl3 Pim1 Pde2a Mrpl23 Psme2 Chordc1 Cebpz B9d2 Gpr65 Pdia6 Manf S100a11 Psme1 Taf1 Gm49602 Pag1 Zfp148 S1pr4 Maff Hivep2 Snhg15 Eif3a Creld2 Otulin Il10 Relb Tnfsf14 Agpat5 Spn Cyba Rfx3 Slc15a3 Efhd2 D1Ertd622e Eef1e1 Usf1 Jund Gimap5 Borcs7 Mapre2 Gm10076 |
| **M0_C** | Hspa1a Hspa1b Utp20 Larp1b Aars Lgals1 Ccl5 Ifitm3 Ssrp1 Emp3 Ifng Myl6 Emb Klrg1 Rap1b Ptms Anxa2 Myo6 Dnaja1 Kcnj8 Pycard Slc29a1 Ptma Ccl3 Clint1 Lgals3 Srm Lsp1 Klra9 Dstn Serf2 Vim Xdh AA467197 Klrb1b Ybx3 Sh3bgrl3 Ormdl3 Icam1 Klra1 Ganc Ccl4 Nfe2l1 Sh2d2a Ubc Timp2 Tyrobp Tomm5 Tm6sf1 |
| **M0_D** | Rbm3 Ctsw Nop58 Txn1 Mybbp1a Hsd11b1 Dkc1 Magohb Nop56 Aprt AI506816 Timm8a1 Ddx18 Nap1l1 Aldoa Las1l Hist1h2be Prdx1 Tomm40 Uqcc2 Park7 Tnfsf12 Klf2 Eif5b Rsl24d1 Itgam mt-Nd4 Abce1 Sec61b Snhg6 Prr13 Pabpc4 Zeb2 Aen Itm2b Dnajc2 Zfas1 Calr Gnb5 Zfp593 Mrps5 Alkbh1 Nars Wdr12 Rpf2 Rrp1b Ythdf2 Larp1 Pdcd11 Eno1 mt-Co2 |

**Table 4**: Top driving genes of single cell trajectories starting in Cluster 2

| Trajectory | cNAÏVE |
|---|---|
| N2_A | S100a6 Lgals1 Myl6 Nkg7 Cd7 Xcl1 Eef1a1 Ppp1r14b Ly6c2 Fgl2 Naca Calr Tcf7 Set Sh3bgrl3 Mrps28 Serf2 mt-Nd2 Phgdh mt-Co1 Atic Mrpl52 mt-Co3 Hsd11b1 Atp5g2 Nhp2 Flna Npm3 Fxyd5 Ltb Itgb2 Ptprcap Klra3 |
| N2_B | Gm47283 Klf3 AI506816 Mdn1 Man1a2 Sbds Fkbp4 Mettl1 Noc2l Selenos Aprt Kcnj8 Pim2 Stmn1 Rrp9 Impdh2 Hnrnpa1 Tuba1b Ugcg Lars Lnpk Cmtm7 Sgk1 Prdx1 Wdr43 Rapgef2 Mcm7 Ndufaf4 Bola2 Ssrp1 Uqcc2 Gadd45g Ypel3 Clock Sms Card19 Nsa2 mt-Nd4l Dut Cnbp |
| N2_C | Khdc1c Coro1a Icam1 Rel Cited2 Prmt3 H2-Q7 Traf1 Cdv3 Tnfsf14 Ep300 Serpina3g Pus7 Kdm2b Nr4a3 Ubald2 Kif20b Vars Lfng Relb Btg2 Lilr4b Gm26917 Pgls Adora2a Dot1l Llph Spred2 Pde2a Trim8 Gpr18 Ppa1 Ms4a4b Eef1e1 Dnmt3a Park7 Tle4 Apbb1ip Gm15472 mt-Nd1 Ubac2 Eef2 Egr3 Itm2b Nfkb2 Pla2g12a Muc1 Pnrc1 Gpr65 Csrnp1 Tgif1 Srpk2 Atg2a |
| N2_D | Hspa1b Fth1 Klf2 Myo6 Klra8 Klra7 Tgfb1 Satb1 Hif1a Nr4a2 Samsn1 Batf Neurl3 Spry2 Ifi203 Klrd1 S100a10 Vps37b Bcl2l11 Pkm 1110038B12Rik Ahnak Cotl1 Eif2s2 Pdcd4 Arid5b Snhg15 Ctsw Slc7a5 Ncr1 Adgre5 Cd53 Tyrobp Ifngr1 Gm19585 mt-Cytb Hsph1 Txn1 Klre1 Ssbp4 Tecpr1 Nolc1 Pml Ifi27 Serinc3 Arid5a Arap2 S100a11 Ssh2 Adap1 Snhg3 Xaf1 |
| **Trajectory** | **cMEMORY** |
| M2_A | Icam1 Pglyrp1 Park7 Slc39a6 Chmp4b Gm10076 Rcc2 Llph Itgam Nfkbiz Cdca7l Eif3a Atic Nifk P2ry14 Ptprcap Larp1 Ifi211 Myc Snhg6 Eno1 Gm26917 Smc1a Lilr4b Klrb1b Polr3d Mrps28 Rap1b Utp11 Asb6 Sptan1 Ezr Wdr12 Pgls |
| M2_B | Hspa1a Hspa1b Eef1g Pa2g4 Aprt Ppa1 Atad3a Impdh2 Eef2 Phf1 Naca Bin2 Cnbp Fau Emb Bzw2 Pabpc4 Lyar Snhg1 Magohb AI506816 Noc2l Apex1 |
| M2_C | Sub1 Nsa2 Ifrd2 Spast Psme2 Crip1 Calm1 Sec61b Anxa2 Plp2 Aen Mrps5 Denr Eif5b Vars Gm20559 Utp18 2810004N23Rik Fkbp2 Klra1 Gars Pdia3 Dnajc2 Lars Ftsj3 Cma1 Gzma Sfxn1 Pycard Aimp1 Xdh Tma16 Rpf2 Hsp90b1 Tars Gadd45gip1 Arhgap45 Kcnj8 Pdcd11 Mcm5 Pus7 |
| M2_D | Ifng Ccl3 Ly6a Furin Ctla2a Gadd45b Nfkbia Fth1 Gzmb Sdc4 Isg15 Myo6 Atf3 AW112010 Srgn Ifit3 Il2ra Tnfrsf8 Hif1a Nr4a2 Tspan13 Zfp36l2 Batf Ms4a4b Neurl3 Spry2 Dnaja1 Ifi203 Calr Klrd1 Bcl2 Rtp4 Vps37b Shisa5 Btg1 Gmfg Junb Otulin Sec61g Tmsb15b1 Herc6 Snhg15 Ctsw Hmgn1 mt-Nd1 Ncr1 Atf6 Rilpl2 Cd53 Ifngr1 mt-Cytb Txn1 Klre1 Rnf213 |
